# Metabolic changes in brain slices over time: a multiplatform metabolomics approach

**DOI:** 10.1101/2020.09.03.280966

**Authors:** Carolina Gonzalez-Riano, Silvia Tapia-González, Gertrudis Perea, Candela González-Arias, Javier DeFelipe, Coral Barbas

## Abstract

Brain slice preparations are widely used for research in neuroscience. However, a high-quality preparation is essential and there is no consensus regarding stable parameters that can be used to define the status of the brain slice preparation after its collection at different time points. Thus, it is critical to establish the best experimental conditions for *ex-vivo* studies using brain slices for electrophysiological recording. In this study, we used a multiplatform (LC-MS and GC-MS) untargeted metabolomics-based approach to shed light on the metabolome and lipidome changes induced by the brain slice preparation process. We have found significant modifications in the levels of 300 compounds, including several lipid classes and their derivatives, as well as metabolites involved in the GABAergic pathway and the TCA cycle. All these preparation-dependent changes in the brain biochemistry should be taken into consideration for future studies to facilitate non-biased interpretations of the experimental results.

## INTRODUCTION

Brain slice preparations are widely used for research in neuroscience for a variety of studies including those focusing on neuronal connectivity, synaptic transmission and plasticity and electrical signaling [1]. Thanks to this approach, fundamental contributions to our understanding of the electrical properties of mammalian neuronal membranes have been achieved [2]. Moreover, diverse therapeutic molecules of novel genes can be efficiently screened by adopting these *ex-vivo* models as a result of the development of disease-relevant slice models that simulate essential features of *in vivo* neurodegenerative pathologies [3]. Recently, electrophysiological studies have begun to be evaluated to examine different aspects of Alzheimer’s disease processes related to different neurophysiological mechanisms underpinning human sensorimotor and cognitive functions. This can be accomplished through the recording of electrophysiological signals from acute brain slices with locally intact neuronal networks and functional connections in specific brain regions [4]. Using these *ex-vivo* preparations, several experimental manipulations can be performed more easily than *in-vivo*, with the additional advantage being that the extracellular environment can be precisely controlled. Indeed, several studies have shown that basic neuronal properties such as firing and local network connectivity found in brain slices are similar to those reported *in vivo* [5].

One major drawback is that the physical and biological traumatic changes induced by *ex-vivo* preparations might change neuronal properties and, therefore, some of the results need to be interpreted cautiously [6]. Indeed, after brain slices’ collection, several alterations that can greatly influence synaptic connectivity have been described, including room temperature, proper oxygenation, and the dissection method [7–10]. Therefore, it is critical to establish the best experimental conditions for the *ex-vivo* studies. However, few reports have tested tissue properties and stability at different time intervals [7]. Pioneering studies focusing on rat hippocampus slices did study such conditions, concluding that brain slices required up to 2.5h to recover from the traumatic dissection process, achieving a steady metabolic and electrophysiological state, regardless of the temperature, glucose, or oxygen levels of slice incubation [11]. Furthermore, a separate study revealed that the general cytoarchitecture was still well-preserved after 6 h of slice incubation [12]. It has also been reported that significant changes in dendritic spine density can be observed in hippocampal slice preparations [13, 14]. However, it has been described that at least 3 h is required for synaptogenesis phenomena taking place in hippocampal slices to promote recovery an ultrastructural state closely resembling the initial *in situ* conditions [15]. Moreover, another critical issue to deal with is the bacterial growth that dramatically reduces the lifespan of the brain slices due to the release of endotoxins such as lipopolysaccharide, which activates the tissue neurodegeneration, affecting cell survival [16]. The addition of antibiotics could hinder bacterial growth. However, these drugs have been suggested as neuronal activators and, hence, could potentially bias results [17]. In summary, there is no consensus regarding stable parameters that can be used to define the status of the tissue after slice collection at different time points. Establishing such a consensus would help researchers to determine appropriate time windows for *ex-vivo* experiments.

Remarkably, brain slicing also induces marked changes in metabolite composition that modify the intracellular and extracellular media, which could affect neural signaling and synaptic activity [18]. Due to the complexity of the brain biochemistry, we have —for the first time— used an integrated untargeted metabolomics strategy employing UHPLC-ESI-QTOF MS and GC-EI-QTOF MS to expand the metabolome coverage, in order to gain a better understanding of the metabolic modifications and the lipid remodeling processes taking place at different time intervals (35 minutes, 2.5 hours, and 5.5 hours) after the collection of brain slices commonly used for electrophysiological recordings.

## METHODS

### Experimental animals and design

This study was performed in 6-week-old male C57BL/6 J mice (Charles River Laboratories, Wilmington, MA). The mice were kept in a 12:12 h light/dark cycle and received food and water ad libitum. All experimental protocols involving the use of animals were performed in accordance with recommendations for the proper care and use of laboratory animals and under the authorization of the regulations and policies governing the care and use of laboratory animals from the Cajal Institute (Madrid, Spain), in accordance with the European Commission (2010/63/EU), FELASA and ARRIVE guidelines. Special care was taken to minimize animal suffering and to reduce the number of animals used to the minimum required for statistical accuracy.

Animals (n=6) were anesthetized with a lethal pentobarbital injection (40 mg/kg BW, Vetoquinol, Madrid, Spain) and transcardially perfused with 10 ml ice-cold (4°C) N-methyl-d-glucamine (NMDG)-HEPES, a modified extracellular solution (92 mM NMDG, 2.5 mM KCl, 1.25 mM NaH_2_PO4, 30 mM NaHCO_3_, 20 mM HEPES, 25 mM glucose, 2 mM thiourea, 5 mM Na-ascorbate, 3 mM Na-pyruvate, 0.5 mM CaCl_2_·2H_2_O, and 10 mM MgSO_4_·7H_2_O), gassed with 95% O_2_/5% CO_2_ pH=7.3[19]. Following decapitation, the mouse brains were rapidly extracted, placed in the same NMDG-HEPES solution (4°C), and cut into coronal sections (350 μm thick) containing hippocampal region using a vibratome (Vibratome VT1200S, Leica). Three consecutive slices were selected from each brain at the level of Bregma −1.22 to −2.70, according to the mouse atlas of Paxinos & Franklin (2001). The slices were then cut through the middle to separate the right and left hemispheres (hemi-slices) and incubated for 10 min at 37°C in NMDG-HEPES solution and gassed with 95% O2/5% CO_2_ (pH = 7.3). This solution preserves the brain slices correctly and improves their viability^1^. The hemi-slices (n=6 per animal) were then transferred to an immersion chamber and perfused with gassed artificial cerebrospinal fluid (ACSF) solution (124 mM NaCl, 2.69 mM KCl, 1.25 mM KH_2_PO_4_, 2 mM MgSO_4_, 26 mM NaHCO_3_, 2 mM CaCl_2_, and 10 mM glucose, gassed with 95% O_2_/5% CO_2_ pH=7.3) at room temperature for the following intervals: 35 min, 2.5h and 5.5h (n=2 animals per interval yielding a total of 12 hemi-slices per interval). The samples were subsequently frozen in liquid nitrogen and stored at −80°C until further analyses to stop any on-going metabolic reactions (Fig. 1).

**Figure 1.**
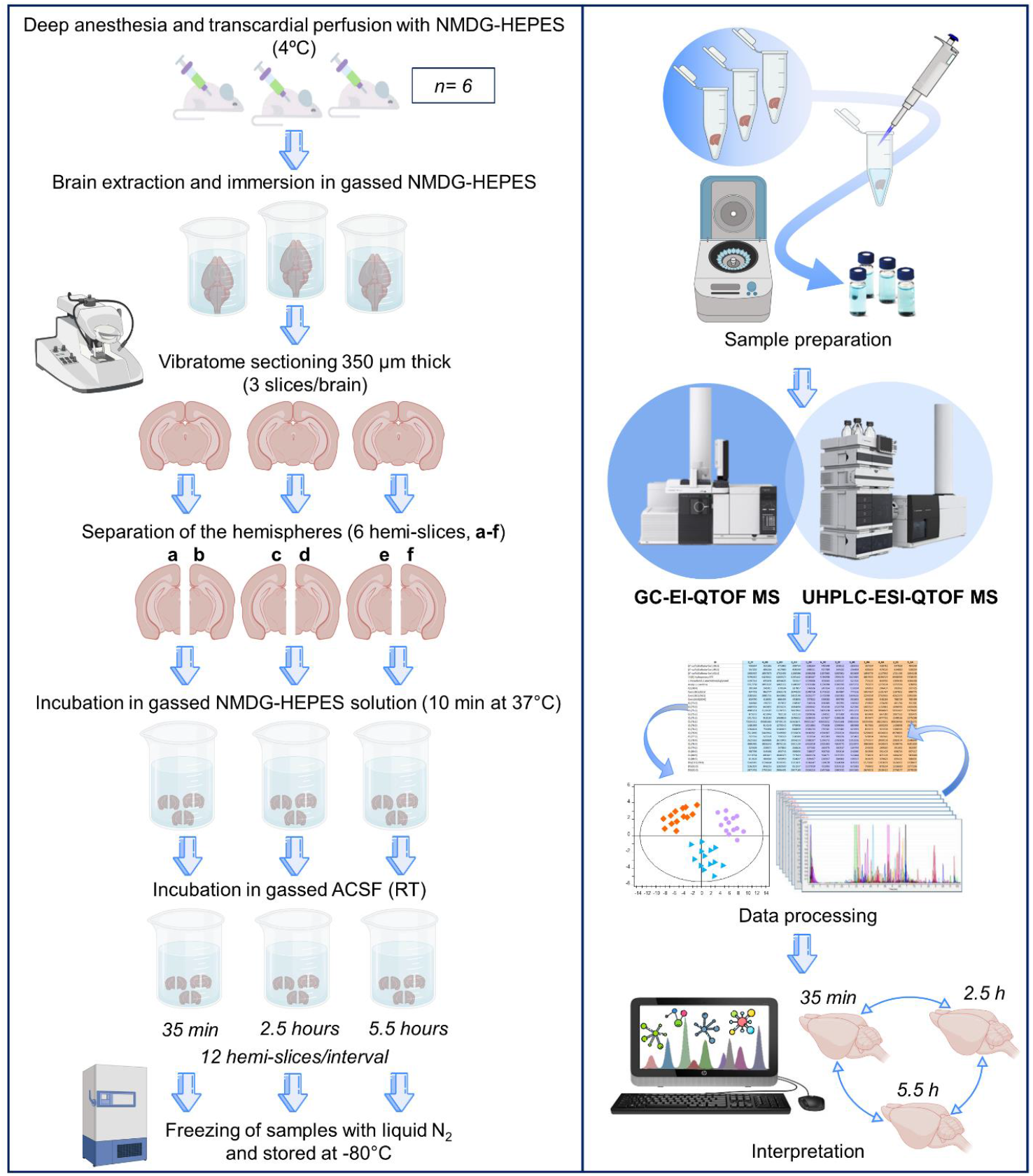
Schematic representation of the experimental design, Vibratome sectioning (350μm thick; 3 slices/brain), brain slice collection, and multiplatform untargeted metabolomics workflow followed in this study.

### Sample processing

The hemi-slices from the 35 min, 2.5h and 5.5h intervals were prepared for GC-MS and LC-MS analyses at CEMBIO (Madrid, Spain). The wet weights of the brain tissue samples ranged from 18.1 to 35.5 mg. The metabolite extraction was performed according to the method previously reported for the analysis of brain tissue [20, 21]. For tissue homogenization, the samples were mixed with cold (−20°C) methanol:water (1:1, v/v) containing 60 ppm of the internal standard 4-nitrobenzoic acid in a ratio of 1 mg tissue:10 μL of solvent per mg of tissue [ratio 1:10 (w/v)]. The disruption of the tissue was achieved using a TissueLyser LT homogenizer (Qiagen, Germany) for 4 minutes. 100 μL of the resulting homogenate was transferred into a 1.5 mL Eppendorf and mixed with 320 μL of cold (−20°C) methanol. The samples were then vortex-mixed for 2 minutes, followed by the addition of 80 μL of methyl tert-butyl ether to extract non-polar metabolites. Subsequently, samples were vortex-mixed for 1 hour at room temperature. After centrifugation at 4000 *g* at 20°C for 20 minutes, 100 μL of each sample was divided into two UHPLC-MS vials with insert (50 μL/each) to inject them directly into the system for LC-MS analyses in positive and negative ionization modes. For GC-MS analysis, 300 μL of supernatant was evaporated to dryness (SpeedVac Concentrator System, Thermo Fisher Scientific, Waltham, MA, USA). The samples were then reconstituted in 20 μL O-methoxyamine hydrochloride (15 mg/mL) in pyridine for methoximation. After vigorous ultrasonication and vortex, the vials were incubated in darkness at room temperature for 16 hours. Next, 20 μL of BSTFA:TMCS (99:1) was added, and the samples were vortex-mixed for 5 min; silylation was carried out for 1 h at 70°C, and finally 100 μL of heptane containing 20 ppm of tricosane was added as an instrumental internal standard (IS).

### Blanks and quality control samples (QCs)

Four blank solutions were prepared along with the rest of the samples, following the same procedure of metabolite extraction (LC-MS and GC-MS) and derivatization (GC-MS). The blank samples were analyzed at the beginning and at the end of the analytical sequence of both techniques. Quality control samples (QCs) were prepared by pooling equal volumes of brain tissue homogenate from each sample and were processed identically in parallel with the rest of the study samples. The samples were then randomized and QCs were injected at the beginning, every 5 experimental samples, and at the end of the batch [22].

### Untargeted metabolomics fingerprinting

The metabolic fingerprinting of brain tissue was performed following a multiplatform untargeted metabolomics-based approach.

#### 1. Brain tissue fingerprinting by UHPLC-ESI-QTOF MS

Metabolomics data from the brain tissue samples were acquired using an Agilent 1290 Infinity II UHPLC system coupled to an Agilent 6545 quadrupole time-of-flight (QTOF) mass spectrometer.

The Agilent 1290 Infinity II Multisampler system equipped with a multiwash option was used to uptake 1.5 μL of samples. To maintain compounds in a stable environment and avoid lipid precipitation, the sampler temperature was kept at 15°C. Reversed-phase chromatography was used in which the mobile phases for positive ionization mode were 5 mM ammonium formate in Milli-Q water (aqueous phase) and 5 mM ammonium formate in methanol:isopropanol (85:15, v/v) (organic phase); whilst for negative ion mode, the mobile phases were 0.1% formic acid in Milli-Q water (aqueous) and 0.1% formic acid in methanol:isopropanol (85:15, v/v), pumped at 0.5 mL/min. The multiwash strategy consisted of a mixture of aqueous phase:organic phase (25:75, v/v), with the wash time set at 30 seconds. An InfinityLab Poroshell 120 EC-C8 (2.1 x 150 mm, 2.7 μm) (Agilent Technologies) column and a suitable guard column (UHPLC Guard 3PK Agilent Technologies) were used and maintained at 60°C. The chromatography gradient started at 75% of organic phase, increasing to 96% at minute 23 and maintained for 8 minutes. The gradient then increased to 100% of organic phase by minute 31.5 and was then maintained for 1 minute until minute 32.5. The starting condition was returned to by minute 33, followed by a 7 min re-equilibration time, taking the total run time to 40 minutes.

The Agilent 6545 QTOF mass spectrometer equipped with a dual AJS ESI ion source was set with the following parameters: 175 V fragmentor, 65V skimmer, 3500 V capillary voltage, 750 V octopole radio frequency voltage, 11 L/min nebulizer gas flow, 290°C gas temperature, 40 psig nebulizer gas pressure, 11 L/min sheath gas flow, 370°C sheath gas temperature. Data were collected in positive and negative ESI modes in separate runs, operated in full scan mode from 100 to 1700 *m/z* with a scan rate of 1.00 spectrum/s. Two reference masses were used over the course of the whole analysis: *m/z* 121.0509 (protonated purine) and *m/z* 922.0098 (protonated HP-921) for the positive ionization mode, and *m/z* 112.9856 (proton-abstracted TFA anion) and *m/z* 966.0007 (formate adduct of HP-921) for the negative mode. These masses were continuously infused into the system at a flow rate of 0.8 mL/min to provide constant mass correction. Auto-MS/MS runs were operated with an MS scan rate of 1 spectrum/s, MS/MS scan rate of 1 spectrum/s, 50–1700 *m/z* mass window, a narrow (~1.3 amu) MS/MS isolation width, and a cycle time of 3.1 s, with collision energy determined on the fly using a slope of 3.6 and intercept of −4.8.

#### 2. Brain tissue fingerprinting by GC-EI-QTOF MS

Data acquisition of brain tissue samples was performed with an Agilent 7890B GC instrument (Agilent Technologies) coupled to a 7250 QTOF mass spectrometer system (Agilent Technologies).

The derivatized sample (1 μL) was injected by an Agilent autosampler (7693) through an Agilent DB5-MS GC Capillary Column (length, 30 m; internal diameter, 0.25 mm; film, 0.25 μm 95% dimethylpolysiloxane/5% diphenylpolysiloxane). The samples were automatically injected in split mode (split ratio 1:12) into a Restek 20782 deactivated glass-wool split liner. The flow rate of helium carrier gas was constant at 0.938 mL/min through the column and the injector temperature was 250°C. The retention time lock (RTL) relative to the internal standard C18:0 methyl ester peak at 19.66 minutes was performed. The temperature gradient was programmed at 60°C (maintained for 1 min), with a ramping rate of 10°C/min up to 325°C. This temperature was then held for 10 minutes before cooling down. The total analysis time was 37.5 min per sample.

The Agilent 7250 QTOF mass spectrometer system equipped with an electron ionization (EI) source was set with the following parameters: filament source temperature, 200°C; detector transfer line temperature, 280°C; quadrupole temperature, 150°C; electron ionization energy, −70 eV. Mass spectra were collected within a mass range of 40–600 m/z at a scan rate of 10 spectra/s.

### Data analysis

#### GC-MS data processing

Once the sequence run had ended, the Total Ion Chromatograms (TIC) obtained were inspected with Agilent MassHunter Qualitative Analysis Navigator (B.08.00), principally to check the quality of the chromatograms and the reproducibility of both of the internal standard signals. Next, raw data files were imported into Agilent Unknowns Analysis Tool (B.09.00) for noise reduction, peak detection, deconvolution, and identification after library search. This software assigned a chemical identity by searching against two targeted libraries: Fiehn library version 2013, and an in-house compound database built with Agilent PCDL Manager (B.08.00) based on the metabolite spectrum and its corresponding retention time (RT), which was experimentally measured after the analysis of the analytical standard. A third commercial library —NIST (National Institute of Standards and Technology, library 2.2 version 2017)— was used in an attempt to identify the unknown compounds. Those metabolites that had a spectrum score ≥ 80% and a concordant retention index based on the *n*-alkane scale were tentatively identified according to this library. The data obtained were then aligned using Agilent MassProfiler Professional (B.15.1) and exported into Agilent MassHunter Quantitative Analysis (B.09.00) for the assignment of the target ions and signal integration. Finally, the data matrix obtained —including the resulting abundances of each metabolite— was normalized according to the IS response (4-nitrobenzoic acid) before any statistical analysis.

#### LC-MS data processing

The data collected after LC-MS analyses in both positive and negative ion modes were reprocessed with Agilent MassHunter Profinder B.08.00 software. The datasets were extracted using the Batch Recursive Feature Extraction (RFE) workflow integrated in the software. This workflow comprises two steps: the Batch Molecular Feature Extraction (MFE) and the Batch Find by Ion feature extraction (FbI). The MFE algorithm consisted of removing unwanted information including the background noise, and then creating a list of possible components that represent the full range of time-of-flight (TOF) mass spectral data features, which are the sum of co-eluting ions that are related by charge-state envelope, isotopologue pattern, and/or the presence of different adducts and dimers. Additionally, the MFE is intended to detect co-eluting adducts of the same feature, selecting the following adducts: [M+H]^+^, [M+Na]^+^, [M+K]^+^ in LC-MS positive ionization; [M–H]^-^, [M+HCOO]^-^, [M+Cl]^-^ in LC-MS negative ion mode, with neutral loss of water also included. The algorithm then aligns the molecular features across the study samples using the mass and RT to build a single spectrum for each compound group. The next step involves FbI using the median values derived from the MFE process to perform a targeted extraction to improve the reliability of finding and reporting features from complex datasets used for differential analysis [23].

### Data normalization and statistical analysis

After obtaining the raw data matrices, an unsupervised Principal Component Analysis (PCA) was performed with SIMCA P+ 15.0 (Umeå, Sweden) to gain an initial overview of the stability and reliability of the metabolomics data before further data analysis. While the QCs were clustered in the GC-MS data, a clear trend in the QCs distribution pattern was observed in the case of the LC-MS analysis corresponding to an intra-batch effect. This intra-batch effect was addressed with a post-acquisition correction based on the QC support vector regression (QC-SVRC), and a radial basis function kernel using MATLAB (R2015a, MathWorks) [24]. Next, the features were selected based on their CV in the QCs; features with CVs over 30% were eliminated.

Finally, differences among the groups were investigated by both univariate (UVDA) and multivariate (MVDA) data analyses. Regarding UVDA, differences among the three stages were evaluated for each individual metabolite using Matlab (R2015a, MathWorks) by one-way ANOVA (p ≤ 0.05) after normality testing applying the Shapiro-Wilk test. Post-hoc, pairwise analyses were performed using the Student’s *t*-test to conclude whether the metabolite was significant or not in a comparison (2.5h *vs* 35min; 5.5h *vs* 35min; 5.5h *vs* 2.5h). Finally, the false discovery rate at level α = 0.05 was checked by Benjamini–Hochberg correction test. Compounds reported with a one-way ANOVA *p* value slightly over 0.05 were kept to enhance the biological information of the study based on their significant *p* value obtained in at least one of the comparisons performed with Student’s *t*-test, and considering that their levels followed the same trend as the metabolites of their class. For MVDA, pareto scaling and logarithmic transformation were applied to the data processing before unsupervised PCA, partial least square-discriminant analysis (PLS-DA), and orthogonal partial least square-discriminant analysis (OPLS-DA) were performed with SIMCA P+ 15.0. The tightness of the clustering of the QCs observed in the PCA plots assessed the reliability and the robustness of the analytical procedure of both platforms. PLS-DA was then performed to reveal the global metabolic changes due to the time interval, and the groups were compared with the OPLS-DA model to maximize class discrimination and investigate the driving forces among the variables. The variable influences on projection (VIP) values were calculated using the OPLS-DA models, keeping those metabolites with a VIP ≥ 1 and a jackknife confidence interval value other than zero. The predictive capacity of the OPLS-DA models generated for each technique was calculated by cross-validation. The data were divided into three parts and each 3^rd^ was then removed in turn. A model was built on the remaining two thirds of the data left in and the randomly selected left-out data was predicted from the new model. This was repeated with each 3^rd^ of the data until all the data had been predicted.

### Compound annotation for LC-MS results

Correctly annotating the metabolites detected by LC-MS analysis is a crucial step to provide a proper biological explanation for the results obtained. A first estimation of the possible metabolite ID was performed by searching the *m/z* of the statistically significant metabolites in the online tool CEU Mass Mediator (http://ceumass.eps.uspceu.es/mediator/) [25, 26]. This search engine comprises the information available in Kegg, HMDB, LIPIDMAPS, Metlin, an in-house library, and MINE. The tentative annotation was based on (i) accurate mass (maximum mass error tolerance 20 ppm); (ii) retention time; (iii) isotopic pattern distribution; (iv) possibility of cation and anion formation; and (v) adduct formation pattern. In parallel, the statistically significant metabolites obtained were extracted in the Auto-MS/MS data. Accurate mass and isotopic distributions for the precursor and product ion were studied for final confirmation of the selected compounds and inspected with Agilent MassHunter Qualitative Analysis Software B.08.00. The metabolites were then identified using different approaches. First, a manual MS/MS spectral interpretation[27] was performed in which a comparison of the MS/MS spectra acquired with available spectral data included in METLIN, MS-DIAL [28], and LIPIDMAPS, together with the software ChemSketch MS Fragmenter (ACD/Laboratories, v.2015.2.5) for product ion structure elucidation, were employed for the assignment of the corresponding identity for each selected metabolite. Once the lipids were annotated, lipidr tool was used for interpretation of the results, based on the lipid classes detected and the structural features of the lipid species in terms of chain length and degree of unsaturation [29].

## RESULTS

We have used the LC-MS and GC-MS analytical platforms to study the metabolic and lipidomic changes taking place at different time intervals after collecting brain samples. The samples were collected and compared at 35 min, 2.5h and 5.5h intervals — and between 2.5h and 5.5h intervals. After an exhaustive analysis, 1182, 878, and 113 features were detected by using LC-MS ESI(+), LC-MS ESI(-), and GC-MS approaches, respectively. After data filtration by coefficient of variation (CV) in quality control (QC) samples (< 30%) and on the basis of the VIP threshold (VIP > 1.0) and *p* ≤ 0.05 in Student’s *t*-test, 165, 113, and 35 metabolites were found to be significant at one time point or more by LC-MS ESI(+), LC-MS ESI(-), and GC-MS analysis, respectively (Supplementary Table 1). The percentage change of each metabolite was also evaluated. The visual inspection of the PCA score plots built for both techniques revealed a tight cluster of the QCs assessing the analytical stability and reproducibility. Additionally, GC-MS plot also revealed a natural grouping of the study samples according to the period elapsed between the tissue collection and the quenching step. The PCA plots and their corresponding explained variance (R^2^) and the predicted variance (Q^2^) are presented in Fig. 2. The PLS-DA analysis showed a clear discrimination among the three groups, with good quality parameters always presenting a difference between R^2^ and Q^2^ lower than 0.3 (Fig. 2). Finally, the OPLS-DA models generated for each analytical method were used to shed light on the most affected metabolites that can be used to determine the principal brain systems altered by the interval time. The quality of the models together with the cross-validation results of each of them are described in Fig. 2. Based on the multivariate and univariate statistical analyses results, critical metabolites involved in the GABAergic system were profoundly affected at different time intervals. During the first period after brain slicing (35min → 2.5h), glutamic acid, α-ketoglutaric acid, pyruvic acid, and fumaric acid levels were upregulated (Fig. 3a,d,f,h). By contrast, glutamine (−42%) and succinic acid (−75%) levels started to decrease 2.5h after collection of brain slices in ACSF solution (Fig. 3b,i). Additionally, GABA, lactic acid and citric acid levels showed a sustained decrease that lasted 5.5h after slicing, confirming the stability of these changes (Fig. 3c,e,g). Similarly, *O*-phosphoethanolamine, aminoadipic acid, and sphingosine also presented such temporal variations (Supplementary Table 1).

**Figure 2.**
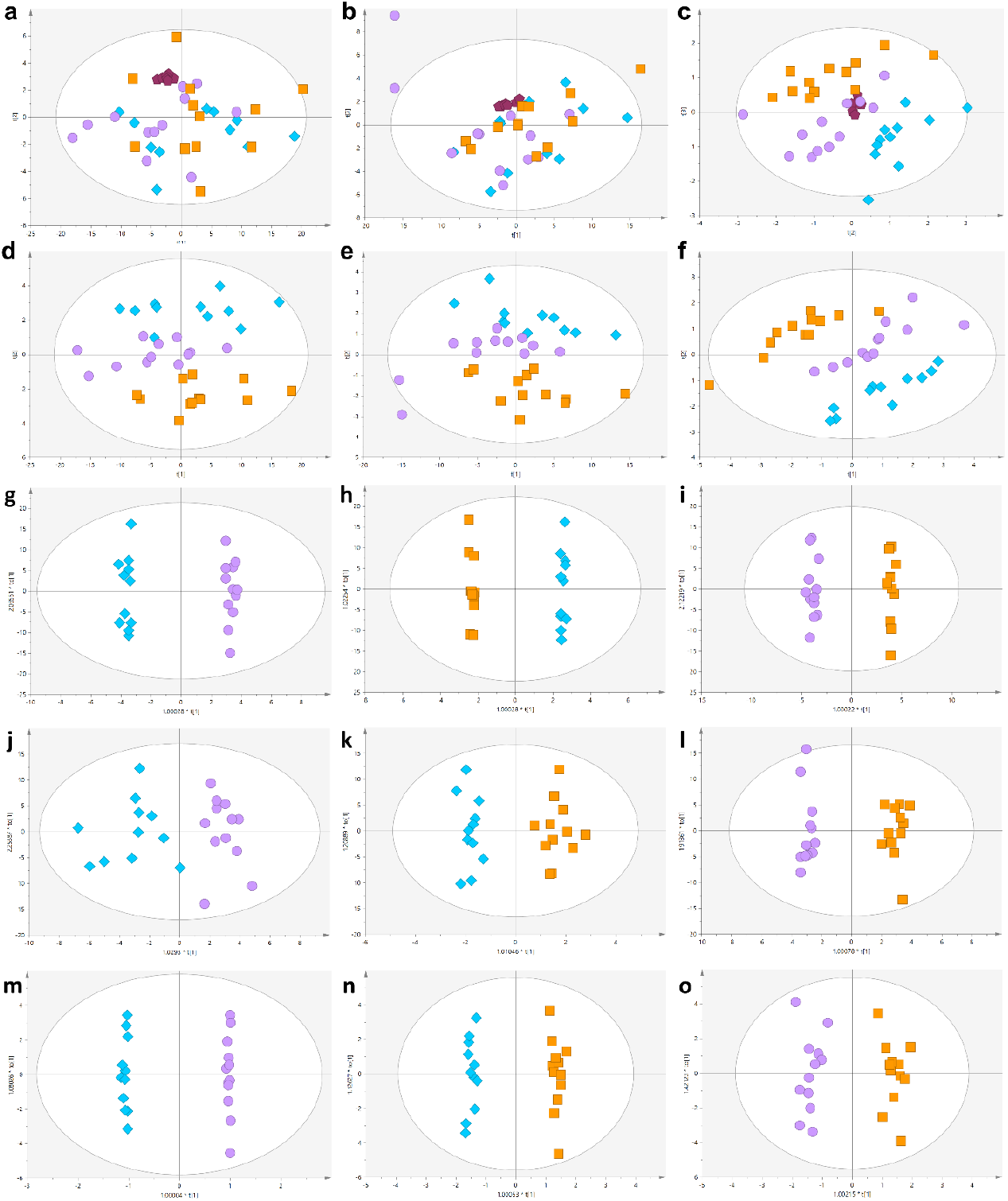
PCA-X and PLS-DA score plots (light blue diamonds, 35min (control samples); purple dots, 2.5h samples; orange squares, 5.5h samples; purple pentagons, QC samples) for the 2 analytical platforms. **a,** (R^2^ = 0.713; Q^2^ = 0.659), **b,** (R^2^ = 0.783; Q^2^ = 0.635), and **c,** (R^2^ = 0.670; Q^2^ = 0.488) represent the PCA-X models for LC-MS ESI (+), LC-MS ESI (-), and GC-MS analyses, respectively. **d,** (R^2^ = 0.527; Q^2^ = 0.379), **e,** (R^2^ = 0.540; Q^2^ = 0.247), and **f,** (R^2^ = 0.921; Q^2^ = 0.749) represent the PLS-DA models built for LC-MS ESI (+), LC-MS ESI (-), and GC-MS analyses, respectively. Supervised OPLS-DA models (light blue diamonds, 35 min (control samples); purple dots, 2.5h samples; orange squares, 5.5h samples)). **g–i,** represent LC-MS ESI(+) data with the following interpretation: 2.5h *vs* 35min R^2^ = 0.994, Q^2^ = 0.822, and percentage of samples correctly classified 95.0% ± 6.9 SD; 5.5h *vs* 35min R^2^ = 0.998, Q^2^ = 0.943, and 95.8% ± 6.5 SD; 5.5h *vs* 2.5h R^2^ = 0.994, Q^2^ = 0.628, and 93.8% ± 7.2 SD. **j–l,** represent LC-MS ESI(-) data with the following interpretation: 2.5h *vs* 35 min R^2^ = 0.804, Q^2^ = 0.136, and 68.8% ± 7.2 SD; 5.5h *vs* 35min R^2^ = 0.943, Q^2^ = 0.622, and 85.0% ± 10.5 SD; 5.5h *vs* 2.5h R^2^ = 0.978, Q^2^ = 0.488, and 75.0% ± 12.5 SD. **m–o,** represent GC-MS data with the following interpretation: 2.5h *vs* 35min R^2^ = 0.999, Q^2^ = 0.827, and 96.4% ± 6.0 SD; 5.5h *vs* 35min R^2^ = 0.990, Q^2^ = 0.956, and 98.2% ± 4.7 SD; 5.5h *vs* 2.5h (R^2^ = 0.955, Q^2^ = 0.737), and 94.6% ± 6.7 SD.

**Figure 3.**
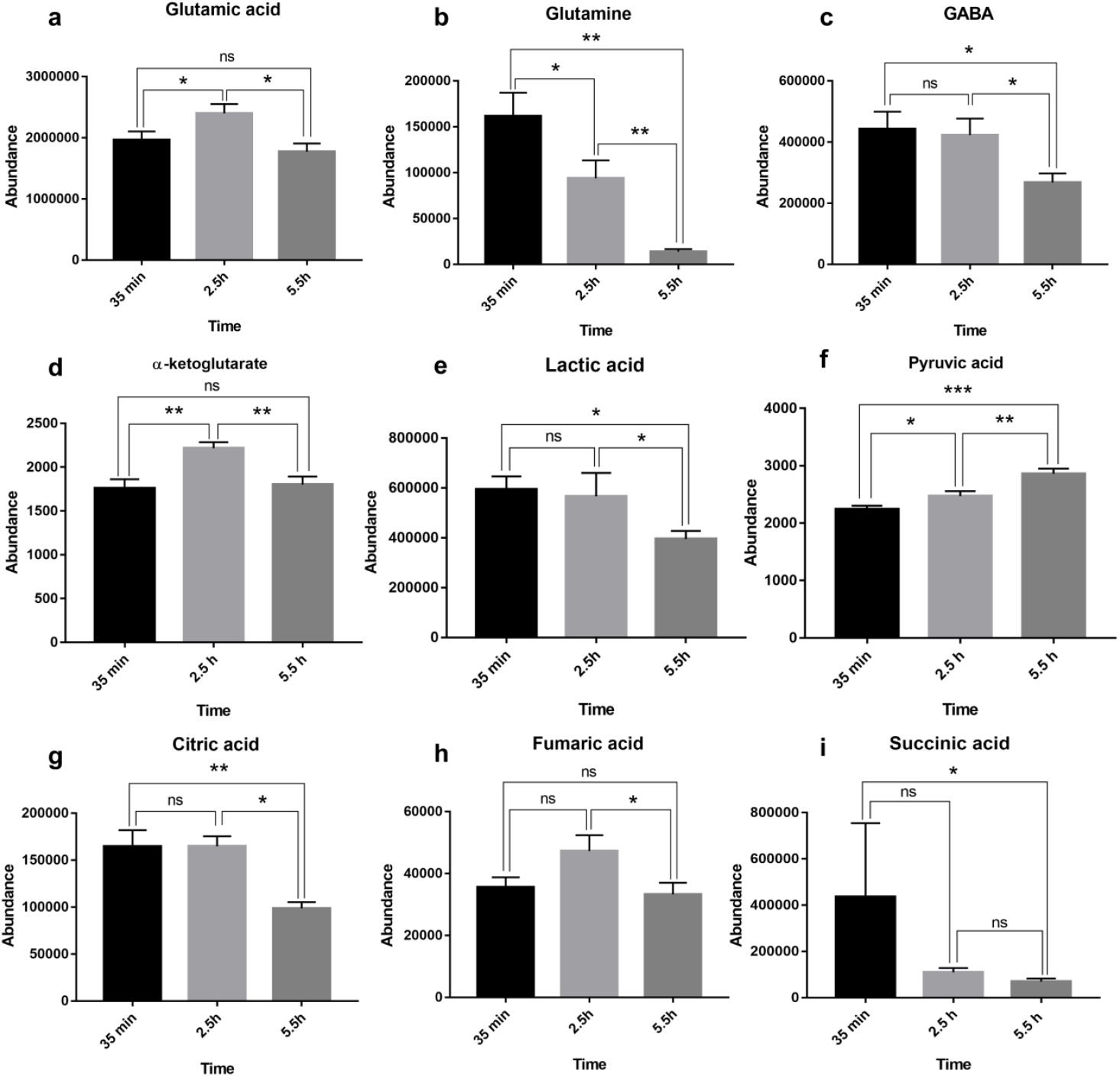
Temporal analysis of the levels of glutamic acid (**a**), glutamine (**b**), GABA, α-ketoglutaric acid (**d**), lactic acid (**e**), pyruvic acid (**f**), citric acid (**g**), fumaric acid (**h**), and succinic acid (**i**). The error bars represent the standard error of the mean (SEM). Student’s *t*-test \**p* value ≤ 0.050; \*\**p* value ≤ 0.010; \*\*\**p* value ≤ 0.001; \*\*\*\**p* value ≤ 0.0001; ns: not significant.

Regarding the lipids, multiple classes were affected by different time intervals including fatty acyls, monoradylglycerols (MAG), diradylglycerols (DAG), triradylglycerols (TAG), glycerophosphocholines (PC), lysophosphocholines (LPC), glycerophosphoethanolamines (PE), lysophosphoethanolamines (LPE), ceramides (Cer), sphingomyelins (SM), among many other classes reported in Supplementary Table 1. However, it should be noted that the changes in lipid levels were not as dramatic as those observed for the amino acids; carboxylic, dicarboxylic and tricarboxylic acids; or the hydroxyl acids (Fig. 4).

**Figure 4.**
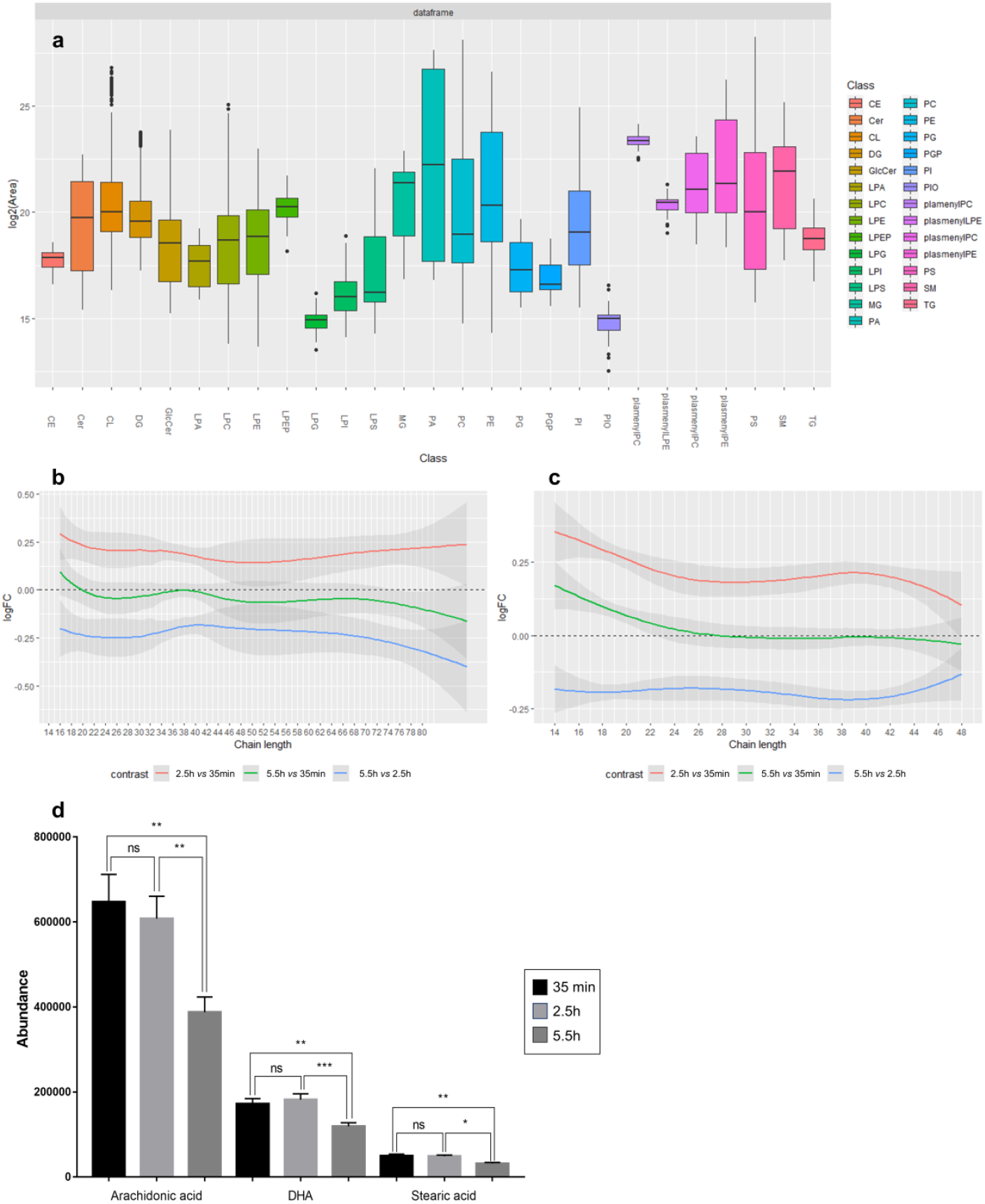
**a,** Intensity distribution boxplots of the principal lipid classes detected as altered by LC-MS analysis. **b**, Lipid chain trend plot of the most relevant lipid species detected by LC-MS ESI(+) and LC-MS ESI(-) (**c**) depending on their carbon atom content. Based on their temporal trend, both plots revealed the tendency for upregulation of lipids after 2.5h of incubation (red), and the downregulation of their levels after 5.5h (blue), nearly reaching their initial levels (green). Graphics generated with lipidr tool. **d,** Temporal variations of arachidonic acid, docosahexaenoic acid, and stearic acid in brain tissue slices. The error bars represent the standard error of the mean (SEM). Student’s *t*-test \**p* value ≤ 0.050; \*\**p* value ≤ 0.010; \*\*\**p* value ≤ 0.001; ns: not significant.

## DISCUSSION

Using a multiplatform untargeted metabolomics-based approach, we performed —for the first time— the metabolic fingerprinting of brain tissue at three time points after brain slicing. We have found significant changes in a large number of compounds, which may serve as a guide for better interpretation of *ex-vivo* experiments (see supplementary material for a list of major compounds analyzed). Since we observed many changes whose possible interpretation is beyond the scope of the present study, what follows is a brief discussion about the possible significance of the changes in the GABAergic pathway, TCA cycle, lipids and lipid derivatives, given their particular relevance in neuronal circuits.

Regarding the GABAergic pathway, 2.5h after brain slicing, there was a clear decrease in critical metabolites of GABAergic synthesis and degradation (**Fig. 5**). Such a reduction of GABA levels would affect the excitation/inhibition balance of brain circuits [30]. Considering the importance of GABAergic control for neuronal excitability, proper information coding and information transfer in the neuronal networks [31], it is crucial to keep the GABAergic tone within the homeostatic range for the physiological excitation/inhibition rate. Accordingly, glutamic acid levels were also decreased 2.5h after slicing, suggesting that homeostatic excitability was achieved and preserved in brain circuits even after the acute dissection process. Although downscaled, this new synaptic signaling would ensure a correct excitation/inhibition balance and favor the study of basic neurophysiological phenomena in *ex-vivo* preparations after 2.5 h of brain slice incubation. Indeed, in brain slices, extensive study has been carried out with regard to neuronal membrane properties and synaptic plasticity processes, such as membrane ion channels and receptors [32], and short- and long-term plasticity [33], which have been later confirmed by *in vivo* studies [34].

**Figure 5.**
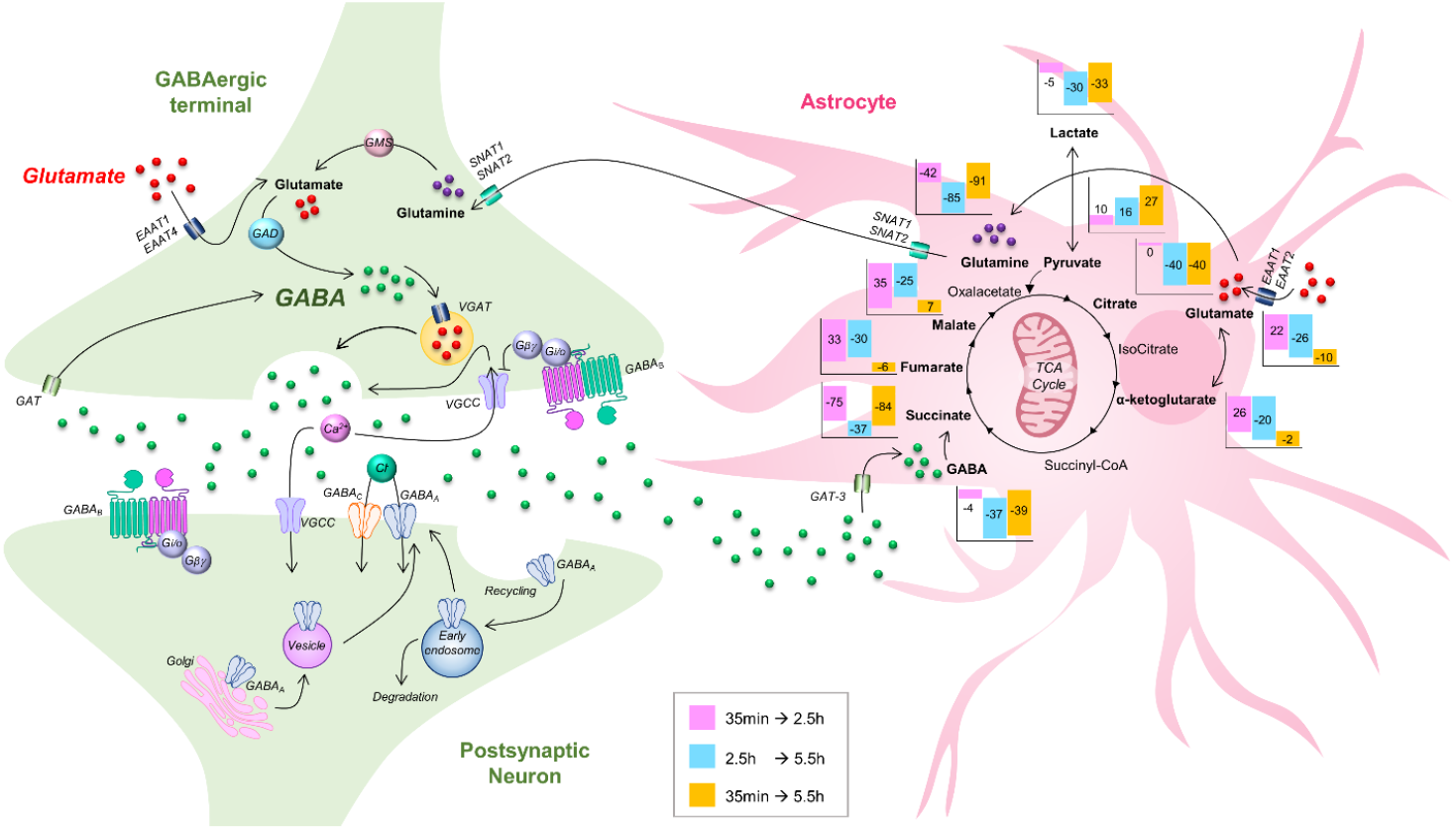
GABA synthesis, release, reuptake and degradation pathway coupled to the TCA cycle. The metabolites in bold were found to be statistically significant for each of the comparisons performed at the different time points. The graphic bars represent the percentage change in the metabolites at each comparison (2.5h *vs* 35 min, pink; 5.5h *vs* 2.5h, blue; 5.5h *vs* 35 min, orange). EAAT1, EAAT2, EAAT4: Excitatory amino acid transporter; GAD: Glutamic acid decarboxylase; GAT: GABA transporter; GAT-3: Glial transporter; GMS: Glutaminase; SNAT1, SNAT2: System A transporter; VGAT: Vesicular GABA transporter; VGCC: Voltage-gated calcium channels.

A well-known hallmark of many neurodegenerative disorders is mitochondrial dysfunction and, consequently, an increment in oxidative stress. The cellular metabolic needs in both physiological and stress conditions can be assumed depending on the state of the mitochondrial networks. Any impairment in the mitochondria will perturb ATP levels and will lead to an imbalance of subsequent reactions involved in numerous functions with catastrophic consequences, ending in cell death [35].

Important changes in the levels of glucose and related metabolites (pyruvate and lactate) have been found 2.5h after slicing. Additionally, many intermediates of the TCA cycle also presented temporary alterations in their levels, including citrate, succinate, fumarate, and malate **(Fig. 3)**. Astrocytes are crucial for brain homeostasis and, in particular, for metabolic support of the brain[36]. Their anatomical position between blood vessels and neurons allows effective glucose uptake from blood capillaries [37]. Once inside astrocytes, glucose can either be stored as glycogen or metabolized to pyruvate [38], which is converted to lactate [39] (Fig. 5). According to the “Astrocyte-to-Neuron Lactate Shuttle” (ANLS) hypothesis [40], lactate is primarily produced by astrocytes and transferred to neurons, where it is converted to pyruvate for aerobic energy production via the TCA cycle [41]. Our results demonstrated that lactate levels started to gradually decrease after 2.5h, while the pyruvate levels started to increase over the same time period (**Fig. 3**). Although there is a lot of evidence supporting the ANLS hypothesis, it remains controversial [42]. Considering that the astrocyte-specific enzymes, pyruvate carboxylase and pyruvate dehydrogenase, can convert two molecules of pyruvate into a new molecule of glutamate [43], different metabolic conditions after brain slicing may indeed challenge cellular signaling pathways in astrocytes, finally affecting both neurons and synaptic activity.

It should be noted that astrocytes efficiently reuptake glutamate released during synaptic events via high-affinity glutamate transporters and contribute to the kinetics of neuronal glutamatergic transmission [44]. The intracellular fate of glutamate in astrocytes may be α-ketoglutarate production [45], or conversion into glutamine [46], which will be released from astrocytes and taken up by neighboring neurons and —after having been metabolized to glutamate— will feed the recycling of synaptic vesicles in the glutamatergic presynaptic terminals [47] (**Fig. 5**). Remarkably, metabolism of glutamate is closely related to GABA, in that glutamate is the immediate precursor of GABA [48]. Thus, the two main excitatory and inhibitory neurotransmitters in the brain are tightly linked, and metabolic alterations that take place in one signaling pathway will affect the other, as has been shown in the present study. Hence, these data indicate that it is not only the neuronal metabolome that is affected by brain slicing, but also glial cells, which contribute to the final state of the circuits (**Fig. 5**).

Lipids and lipid derivatives play a vital role in the correct neuronal function in the brain, representing up to 50% of the brain dry/wet ratio. The lipid landscape present in brain tissue across its areas, cell types, and cell compartments is extraordinarily diverse and complex. Glycerophospholipids (GPL) are highly involved in signal transduction and neurodegeneration, and they are the most abundant lipid category in adult brain. We found that the principal fractions of lipids affected were glycerophosphocholine (PC), glycerophosphoethanolamine (PE), glycerophosphoinositol (PI), glycerophosphoserine (PS), and their corresponding lyso-forms (**Fig. 4a**). The levels of all GPL species showed a significant increase 2.5h after brain tissue collection, followed by a sharp decrease, nearly reaching their initial levels after 5.5h (**Fig. 4b–c**). Indeed, the levels of sn-glycero-3-phosphoethanolamine, which is involved in PE synthesis, started to decrease after 2.5h and remained low after 5.5h — suggesting that it had been ‘consumed’ for this purpose (i.e., the synthesis of PE). Sphingolipids, which are structural components of the membranes, are also involved in many vital functions for correct brain functioning and development, including cellular regulation, neuronal differentiation, synaptic transmission, and myelin sheath stabilization [49]. In this regard, sphingolipids —including ceramides, sphingomyelins, and neutral and acidic glycosphingolipids— displayed a similar trend to that of GPLs. The brain slicing itself is a disruptive procedure that shatters the cell membranes through the slicing region. Therefore, the viable cells that remain confined between two layers of damaged tissue start to seal their membranes in order to avoid the spillover of cellular content originating from the adjacent damaged tissue [50]. Consequently, it can be hypothesized that the increase in the levels of all GPLs and sphingolipid classes found within 2.5h after brain slicing may be related to an upregulation in lipid biosynthesis inducing cell membrane rearrangements to reduce cellular leakage, followed by a stabilization of the tissue within the period between 2.5 and 5.5h after slicing. Additionally, after 5.5h of brain slice incubation, a decrease in serine levels was observed (Supplementary Table 1). Given that serine is essential for the synthesis of both sphingolipids and phosphatidylserines, the decrease in its levels suggested that it had been consumed for this purpose. Interestingly, D-Serine, which is reportedly synthetized and released by astrocytes [51], is a co-agonist of NMDA receptors [52]. Therefore, the above-mentioned serine reduction would affect NMDA-dependent glutamatergic signaling in brain circuits after sectioning, which — in parallel with GABAergic metabolite downregulation— would contribute to homeostatic synaptic balance.

Regarding the omega-6 PUFA arachidonic acid (ARA, 20:4n-6) and the omega-3 PUFA docosahexaenoic acid (DHA, 22:6n-3), these two acids make up approximately 20% of the free fatty acid content in mammalian brain tissue [53]. Both lipid species are involved in several key aspects of brain function (either by themselves or interacting with other PUFAs) —including brain metabolism and structure. ARA is a cell diffusible fatty acid that is thought to serve as an intercellular messenger in several parts of the CNS, controlling the electrical and biochemical behavior of neurons and glial cells, and it is also the precursor of a large number of bioactive eicosanoid products [54]. Additionally, the omega-3 PUFAs can modulate sodium and calcium channels in the hippocampal CA1 neurons, prolonging the inactivated state of these channels [55]. The main consequence of the electrophysiological effect of these fatty acids is a anticonvulsant action in the CNS [56]. Although there is a balance between the ARA and DHA content in brain tissue, both lipids present differences regarding their GPL location within the brain since ARA greatly exceeds DHA in PI lipids, whereas DHA exceeds ARA in PS lipids [57]. Interestingly, our results showed that 35% of the PI species affected contained ARA, while 22% of the altered PS species contained DHA (Supplementary Table 1). The storage and biosynthesis of PS in neuronal tissues depends on the levels of membrane DHA and stearic acid (18:0). Moreover, 20% of the brain’s total energy is provided by the mitochondrial β-oxidation of fatty acids. Our findings revealed that the levels of ARA, DHA and stearic acid remained stable for up to 2.5h of incubation and then dramatically decreased (**Fig. 4d**).

Oleamide is a fatty acid amide that can be synthesized *de novo* within the brain microsomes using oleic and ammonia as substrates. It belongs to the endocannabinoid family and may act as a sleep-inducing agent [58]. This endogenous lipid mediator is elevated in the central nervous system of sleep-deprived mammals, probably via its direct agonist action on CB1 cannabinoid receptors. Moreover, oleamide interacts with voltage-gated Na^+^ channels and allosterically with GABA_A_ and 5-HT receptors [59]. We detected a large and significant increment in the oleamide levels after 2.5h of incubation, maintaining these high levels after 5.5h of incubation (Supplementary Table 1), which would affect the synaptic events mediated by these receptors.

## CONCLUSIONS

In summary, we have shown —for the first time in a preparation used for electrophysiological recordings— that complex changes in the metabolome and lipidome of brain slices take place in the cells after the brain slicing process and we have highlighted the potential impact that these metabolites would have on *ex-vivo* studies. Our findings from a multiplatform untargeted metabolomics experiment unveiled modifications in the levels of 300 compounds, including amino acids; carboxylic and dicarboxylic acids; hydroxy acids; and multiple lipid classes and their derivatives. In addition, we can conclude that at least 5.5h is required for the brain lipidome to recover after collection of the slices, highlighting the impact that the brain slicing procedure has on the brain structure. All these preparation-dependent changes in the brain metabolome should be taken into consideration, since knowledge about which metabolites are stable and which are susceptible to change —as well as information about their potential effects on electrophysiological recordings— is essential to make non-biased interpretations about the experimental results.

## Abbreviations

ARA: arachidonic acid
Cer: ceramides
CV: coefficient of variation
DAG: diradylglycerols
DHA: docosahexaenoic acid
ESI: electrospray ionization
FbI: find by ion
GABA: gammaAminobutyric acid
GC-MS: gas chromatography-mass spectrometry
GPLs: glycerophospholipids
h: hour
HMDB: human metabolome database
IS: internal standard
LC-MS: liquid chromatography-mass spectrometry
LPC: lysoglycerophosphocholines
LPE: lysoglycerophosphoethanolamines
MAG: monoradylglycerols
MFE: molecular feature extraction
min: minutes
MVDA: multivariate data analysis
NIST: National Institute of Standards and Technology
NMDA: N-methyl-D-aspartate
NMDG: N-methyl-d-glucamine
ns: not significant
OPLS-DA: orthogonal partial least square-discriminant analysis
PC: glycerophosphocholines
PCA: principal component analysis
PE: glycerophosphoethanolamines
PI: glycerophosphoinositols
PLS-DA: partial least square-discriminant analysis
PS: glycerophosphoserine
PUFA: polyunsaturated fatty acid
QCs: quality controls
QC-SVRC: quality control-support vector regression
QTOF: quadrupole time-of-flight
RFE: recursive feature extraction
RT: retention time
RTL: retention time lock
SD: standard deviation
SEM: standard error of mean
SM: sphingomyelins
TAG: triradylglycerols
TIC: total ion chromatogram
TCA: tricarboxylic acid cycle
UVDA: univariate data analysis
VIP: variable influence on projection

## DECLARATIONS

### Ethics approval and consent to participate

All experimental protocols involving the use of animals were performed in accordance with recommendations for the proper care and use of laboratory animals and under the authorization of the regulations and policies governing the care and use of laboratory animals from the Cajal Institute (Madrid, Spain), in accordance with the European Commission (2010/63/EU), FELASA and ARRIVE guidelines. Special care was taken to minimize animal suffering and to reduce the number of animals used to the minimum required for statistical accuracy.

### Consent for publication

Not applicable.

### Availability of data and materials

The data that support the findings of this study are available from the corresponding authors upon request.

### Author contributions

Conceptualization: Silvia Tapia-González, Gertrudis Perea, Javier DeFelipe.

Data acquisition: Carolina Gonzalez-Riaño, Coral Barbas.

Data analysis: Carolina Gonzalez-Riaño, Coral Barbas.

Data interpretation: Carolina Gonzalez-Riaño, Gertrudis Perea, Silvia Tapia-González, Candela González-Arias, Javier DeFelipe, Coral Barbas.

Supervision: Coral Barbas, Javier DeFelipe.

Writing – original draft: Carolina Gonzalez-Riaño, Gertrudis Perea, Javier DeFelipe.

Writing – review & editing: Carolina Gonzalez-Riaño, Gertrudis Perea, Silvia Tapia-González, Candela González-Arias, Javier DeFelipe, Coral Barbas.

## Acknowledgements

We would like to thank Nick Guthrie for his excellent text editing, and Vanesa Alonso for her technical assistance.

## SUPPLEMENTARY INFORMATION

### Additional file

Additional file 1: **Supplementary Methods.** Table 1 – Metabolites found as statistically significant at any of the comparisons performed at different time points.

## Supplementary information

**Supplementary Table 1.**
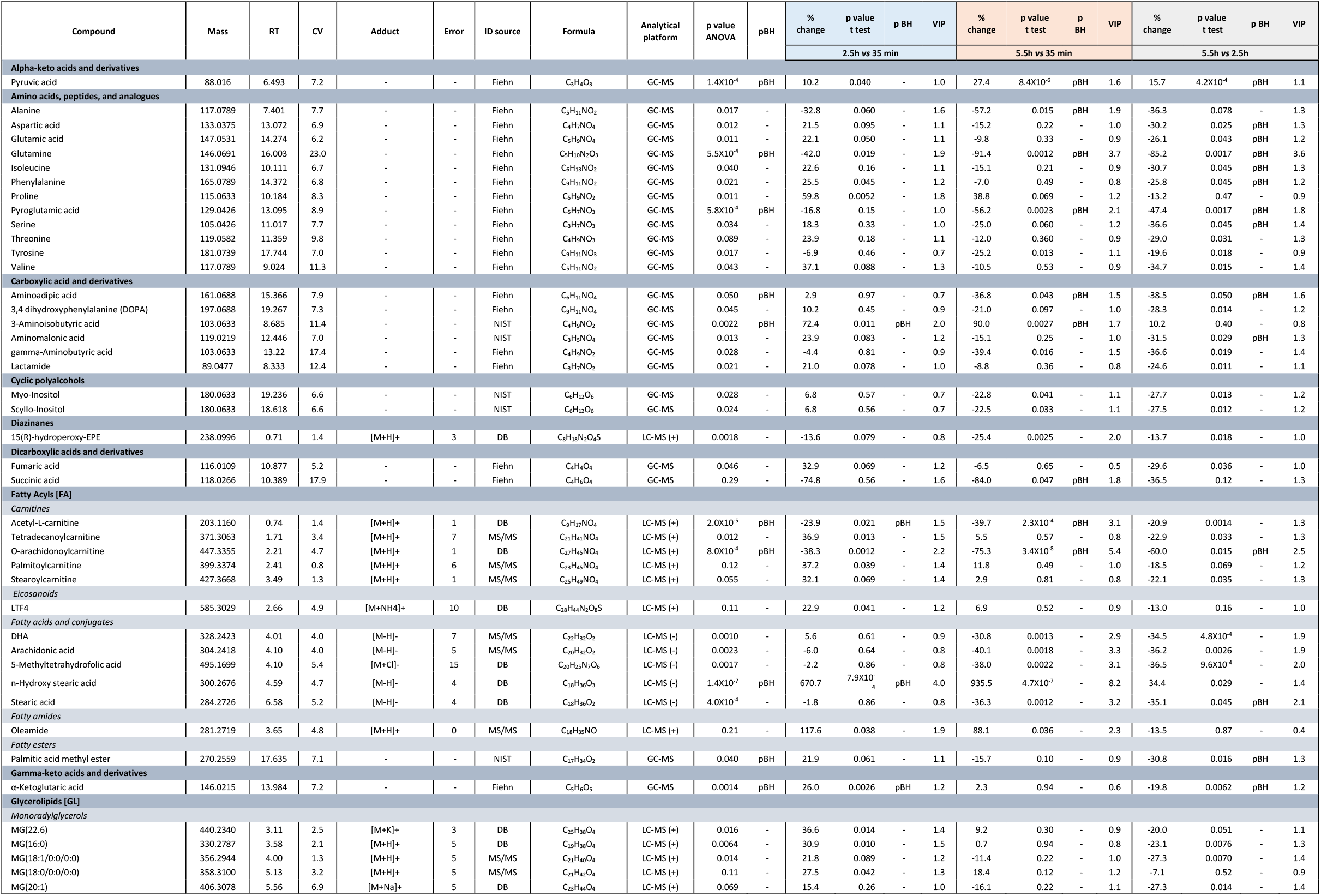
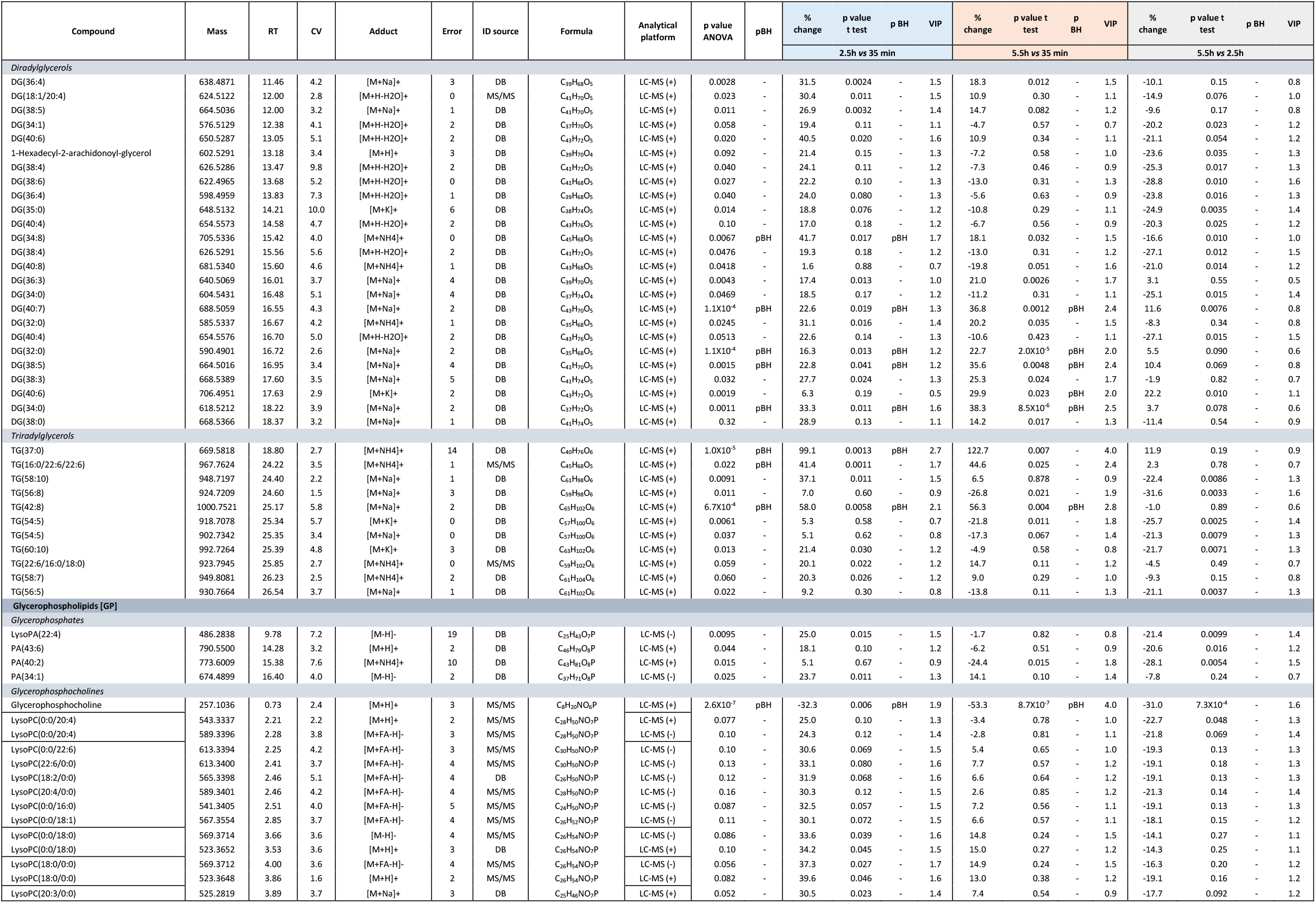
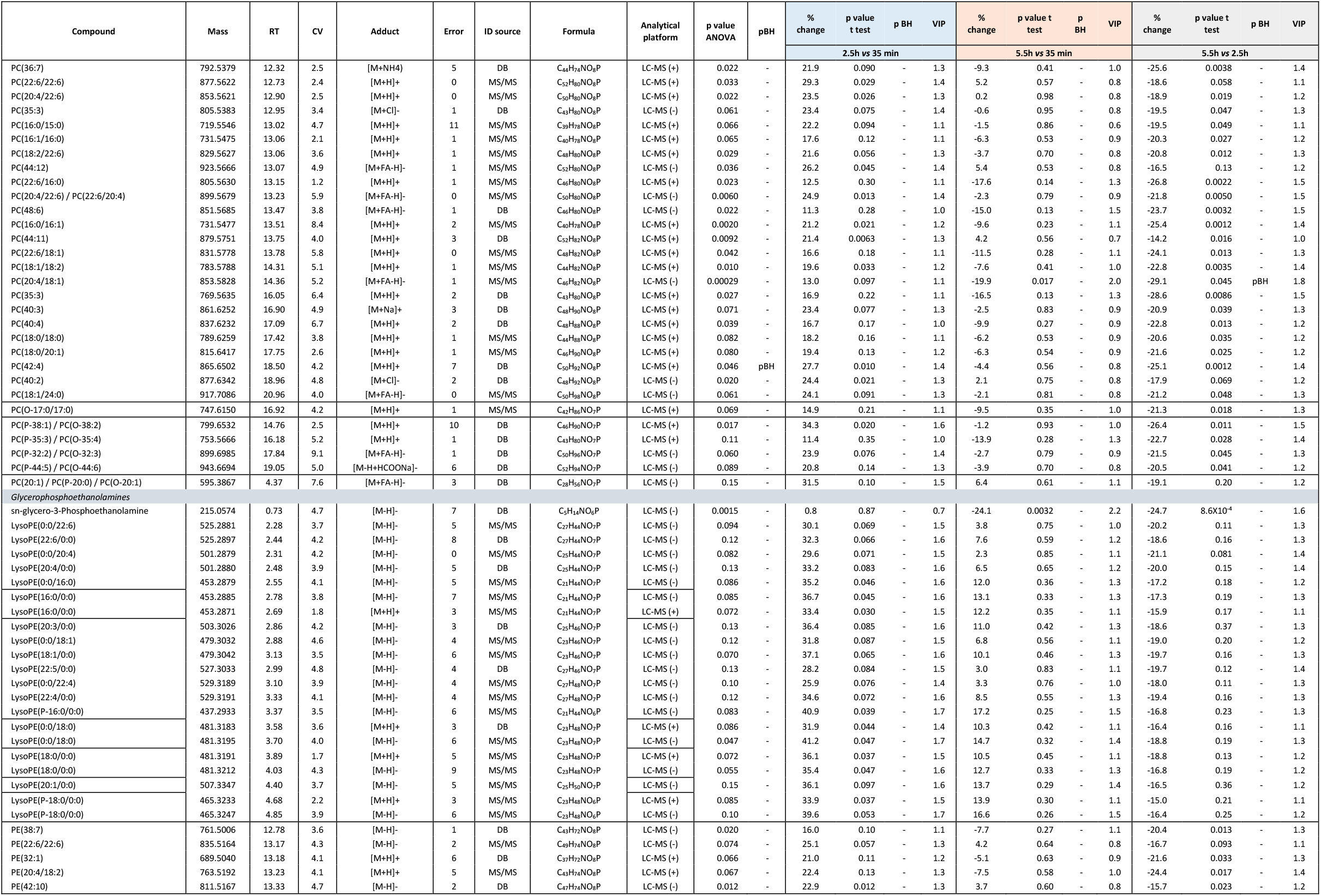
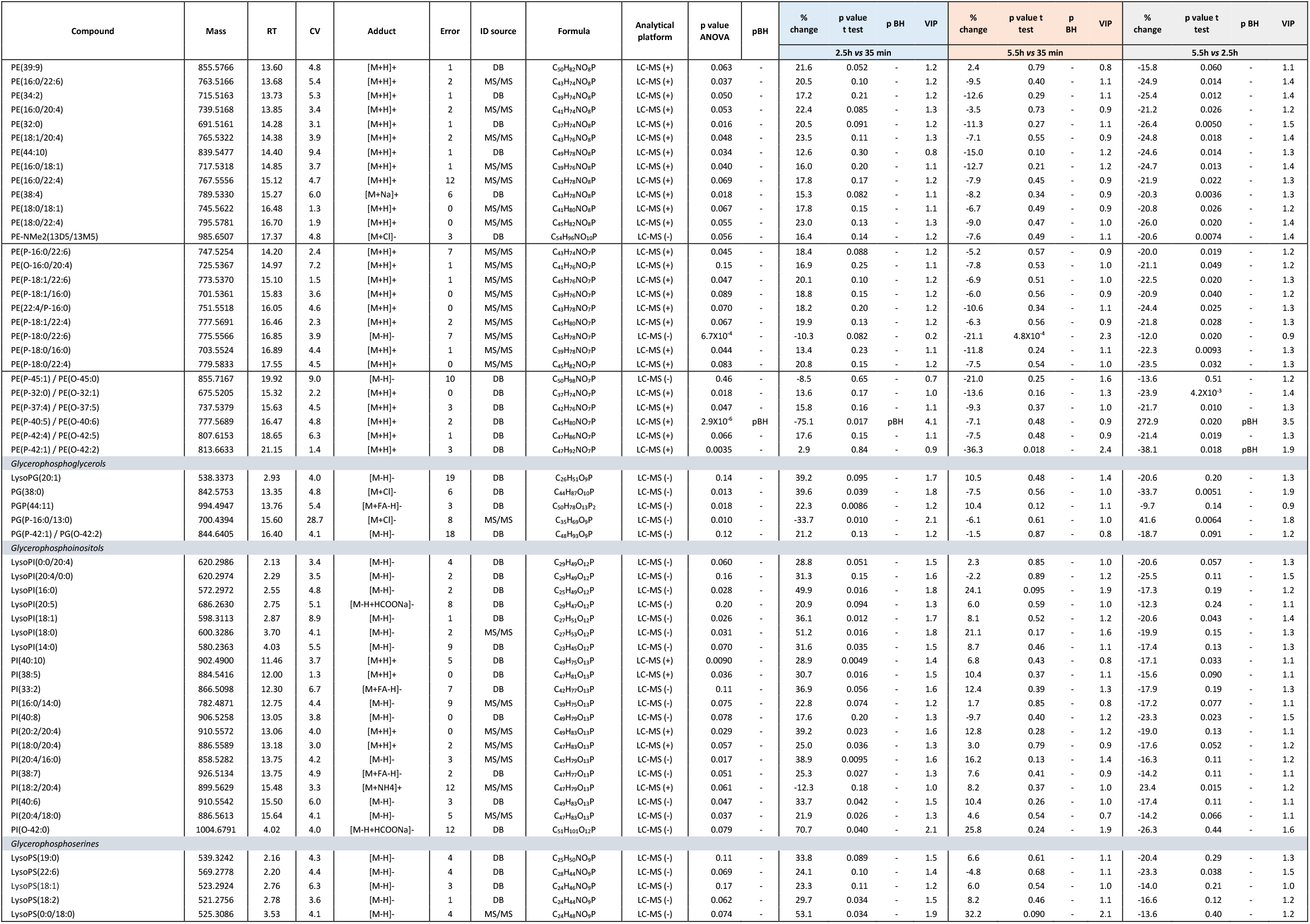
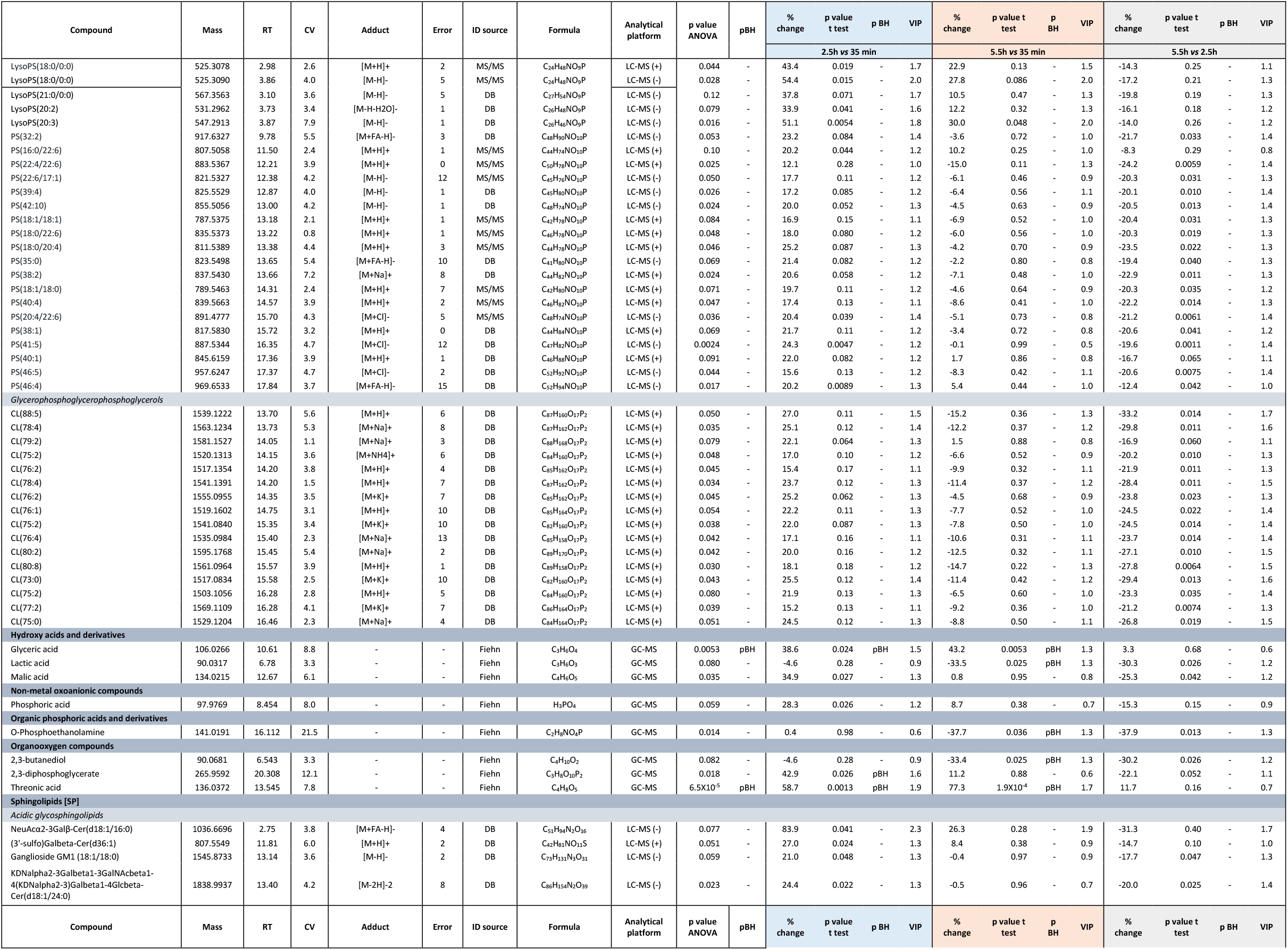
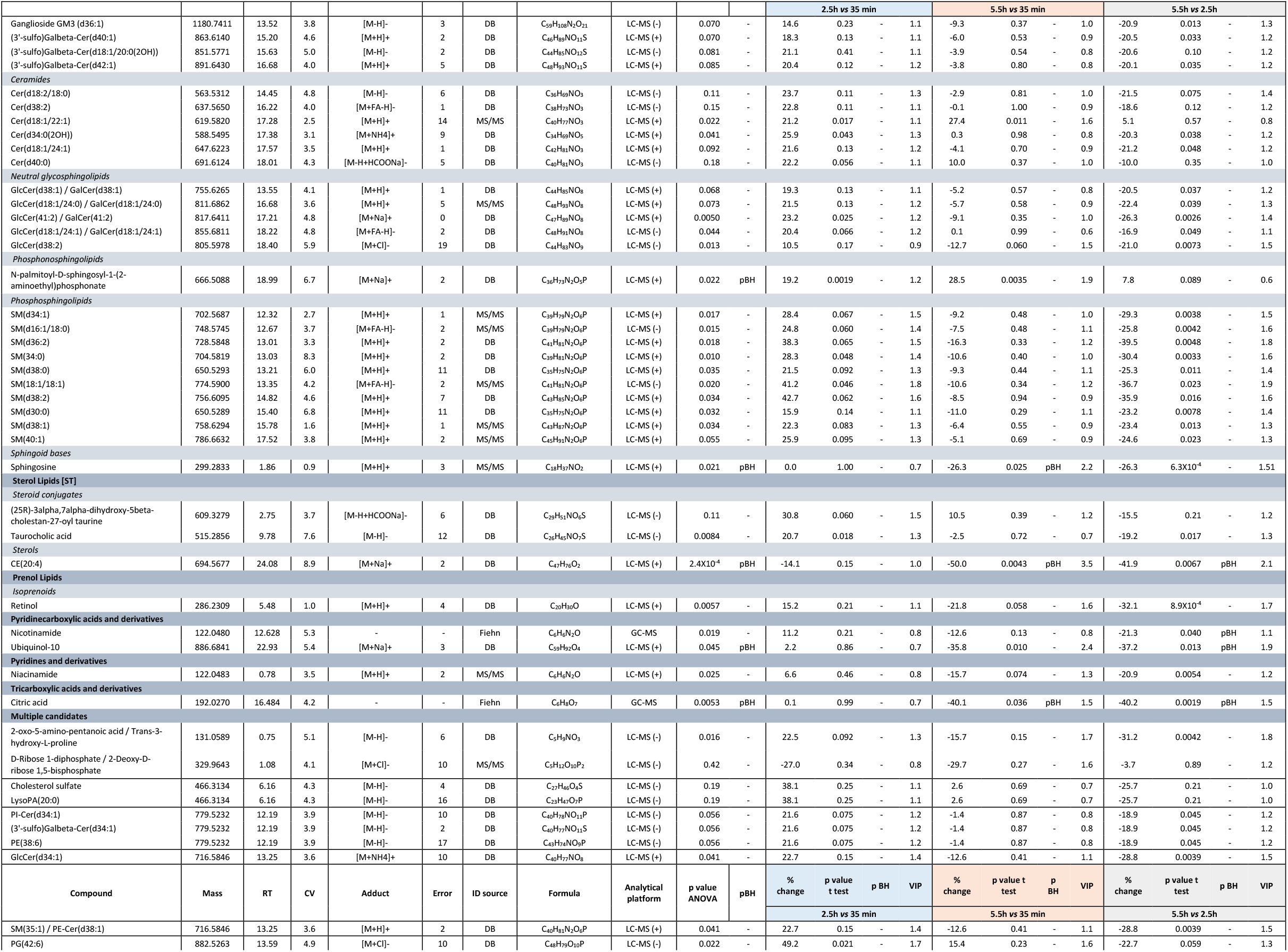
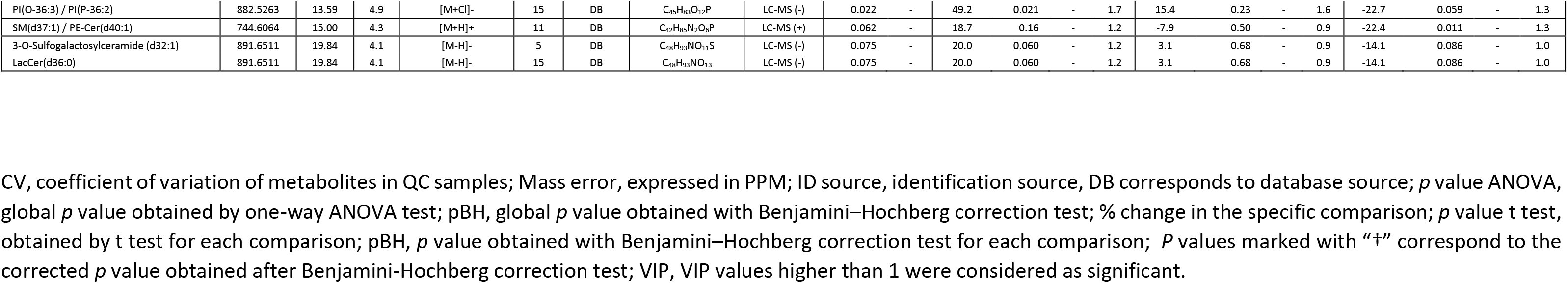
Metabolites found as statistically significant at any of the comparisons performed at different time points.

