## Supplementary Table 1 for "Metabolic changes in brain slices over time: a multiplatform metabolomics approach"

Metabolites found as statistically significant at any of the comparisons performed at different time points.

| Compound | Mass | RT | CV | Adduct | Error | ID source | Formula | Analytical platform | p value ANOVA | pBH | % change | p value t test | p BH | VIP | % change | p value t test | p BH | VIP | % change | p value t test | p BH | VIP |
|---|---|---|---|---|---|---|---|---|---|---|---|---|---|---|---|---|---|---|---|---|---|---|
| | | | | | | | | | | | 2.5h *vs* 35 min | | | | 5.5h *vs* 35 min | | | | 5.5h *vs* 2.5h | | | |
| **Alpha-keto acids and derivatives** | | | | | | | | | | | | | | | | | | | | | | |
| Pyruvic acid | 88.016 | 6.493 | 7.2 | - | - | Fiehn | $C_3H_4O_3$ | GC-MS | $1.4X10^{-4}$ | pBH | 10.2 | 0.040 | - | 1.0 | 27.4 | $8.4X10^{-6}$ | pBH | 1.6 | 15.7 | $4.2X10^{-4}$ | pBH | 1.1 |
| **Amino acids, peptides, and analogues** | | | | | | | | | | | | | | | | | | | | | | |
| Alanine | 117.0789 | 7.401 | 7.7 | - | - | Fiehn | $C_5H_{11}NO_2$ | GC-MS | 0.017 | - | -32.8 | 0.060 | - | 1.6 | -57.2 | 0.015 | pBH | 1.9 | -36.3 | 0.078 | - | 1.3 |
| Aspartic acid | 133.0375 | 13.072 | 6.9 | - | - | Fiehn | $C_4H_7NO_4$ | GC-MS | 0.012 | - | 21.5 | 0.095 | - | 1.1 | -15.2 | 0.22 | - | 1.0 | -30.2 | 0.025 | pBH | 1.3 |
| Glutamic acid | 147.0531 | 14.274 | 6.2 | - | - | Fiehn | $C_5H_9NO_4$ | GC-MS | 0.011 | - | 22.1 | 0.050 | - | 1.1 | -9.8 | 0.33 | - | 0.9 | -26.1 | 0.043 | pBH | 1.2 |
| Glutamine | 146.0691 | 16.003 | 23.0 | - | - | Fiehn | $C_5H_{10}N_2O_3$ | GC-MS | $5.5X10^{-4}$ | pBH | -42.0 | 0.019 | - | 1.9 | -91.4 | 0.0012 | pBH | 3.7 | -85.2 | 0.0017 | pBH | 3.6 |
| Isoleucine | 131.0946 | 10.111 | 6.7 | - | - | Fiehn | $C_6H_{13}NO_2$ | GC-MS | 0.040 | - | 22.6 | 0.16 | - | 1.1 | -15.1 | 0.21 | - | 0.9 | -30.7 | 0.045 | pBH | 1.3 |
| Phenylalanine | 165.0789 | 14.372 | 6.8 | - | - | Fiehn | $C_9H_{11}NO_2$ | GC-MS | 0.021 | - | 25.5 | 0.045 | - | 1.2 | -7.0 | 0.49 | - | 0.8 | -25.8 | 0.045 | pBH | 1.2 |
| Proline | 115.0633 | 10.184 | 8.3 | - | - | Fiehn | $C_5H_9NO_2$ | GC-MS | 0.011 | - | 59.8 | 0.0052 | - | 1.8 | 38.8 | 0.069 | - | 1.2 | -13.2 | 0.47 | - | 0.9 |
| Pyroglutamic acid | 129.0426 | 13.095 | 8.9 | - | - | Fiehn | $C_5H_7NO_3$ | GC-MS | $5.8X10^{-4}$ | pBH | -16.8 | 0.15 | - | 1.0 | -56.2 | 0.0023 | pBH | 2.1 | -47.4 | 0.0017 | pBH | 1.8 |
| Serine | 105.0426 | 11.017 | 7.7 | - | - | Fiehn | $C_3H_7NO_3$ | GC-MS | 0.034 | - | 18.3 | 0.33 | - | 1.0 | -25.0 | 0.060 | - | 1.2 | -36.6 | 0.045 | pBH | 1.4 |
| Threonine | 119.0582 | 11.359 | 9.8 | - | - | Fiehn | $C_4H_9NO_3$ | GC-MS | 0.089 | - | 23.9 | 0.18 | - | 1.1 | -12.0 | 0.360 | - | 0.9 | -29.0 | 0.031 | - | 1.3 |
| Tyrosine | 181.0739 | 17.744 | 7.0 | - | - | Fiehn | $C_9H_{11}NO_3$ | GC-MS | 0.017 | - | -6.9 | 0.46 | - | 0.7 | -25.2 | 0.013 | - | 1.1 | -19.6 | 0.018 | - | 0.9 |
| Valine | 117.0789 | 9.024 | 11.3 | - | - | Fiehn | $C_5H_{11}NO_2$ | GC-MS | 0.043 | - | 37.1 | 0.088 | - | 1.3 | -10.5 | 0.53 | - | 0.9 | -34.7 | 0.015 | - | 1.4 |
| **Carboxylic acid and derivatives** | | | | | | | | | | | | | | | | | | | | | | |
| Aminoadipic acid | 161.0688 | 15.366 | 7.9 | - | - | Fiehn | $C_6H_{11}NO_4$ | GC-MS | 0.050 | pBH | 2.9 | 0.97 | - | 0.7 | -36.8 | 0.043 | pBH | 1.5 | -38.5 | 0.050 | pBH | 1.6 |
| 3,4 dihydroxyphenylalanine (DOPA) | 197.0688 | 19.267 | 7.3 | - | - | Fiehn | $C_9H_{11}NO_4$ | GC-MS | 0.045 | - | 10.2 | 0.45 | - | 0.9 | -21.0 | 0.097 | - | 1.0 | -28.3 | 0.014 | - | 1.2 |
| 3-Aminoisobutyric acid | 103.0633 | 8.685 | 11.4 | - | - | NIST | $C_4H_9NO_2$ | GC-MS | 0.0022 | pBH | 72.4 | 0.011 | pBH | 2.0 | 90.0 | 0.0027 | pBH | 1.7 | 10.2 | 0.40 | - | 0.8 |
| Aminomalonic acid | 119.0219 | 12.446 | 7.0 | - | - | NIST | $C_3H_5NO_4$ | GC-MS | 0.013 | - | 23.9 | 0.083 | - | 1.2 | -15.1 | 0.25 | - | 1.0 | -31.5 | 0.029 | pBH | 1.3 |
| gamma-Aminobutyric acid | 103.0633 | 13.22 | 17.4 | - | - | Fiehn | $C_4H_9NO_2$ | GC-MS | 0.028 | - | -4.4 | 0.81 | - | 0.9 | -39.4 | 0.016 | - | 1.5 | -36.6 | 0.019 | - | 1.4 |
| Lactamide | 89.0477 | 8.333 | 12.4 | - | - | Fiehn | $C_3H_7NO_2$ | GC-MS | 0.021 | - | 21.0 | 0.078 | - | 1.0 | -8.8 | 0.36 | - | 0.8 | -24.6 | 0.011 | - | 1.1 |
| **Cyclic polyalcohols** | | | | | | | | | | | | | | | | | | | | | | |
| Myo-Inositol | 180.0633 | 19.236 | 6.6 | - | - | NIST | $C_6H_{12}O_6$ | GC-MS | 0.028 | - | 6.8 | 0.57 | - | 0.7 | -22.8 | 0.041 | - | 1.1 | -27.7 | 0.013 | - | 1.2 |
| Scyllo-Inositol | 180.0633 | 18.618 | 6.6 | - | - | NIST | $C_6H_{12}O_6$ | GC-MS | 0.024 | - | 6.8 | 0.56 | - | 0.7 | -22.5 | 0.033 | - | 1.1 | -27.5 | 0.012 | - | 1.2 |
| **Diazinanes** | | | | | | | | | | | | | | | | | | | | | | |
| 15(R)-hydroperoxy-EPE | 238.0996 | 0.71 | 1.4 | [M+H]+ | 3 | DB | $C_8H_{18}N_2O_4S$ | LC-MS (+) | 0.0018 | - | -13.6 | 0.079 | - | 0.8 | -25.4 | 0.0025 | - | 2.0 | -13.7 | 0.018 | - | 1.0 |
| **Dicarboxylic acids and derivatives** | | | | | | | | | | | | | | | | | | | | | | |
| Fumaric acid | 116.0109 | 10.877 | 5.2 | - | - | Fiehn | $C_4H_4O_4$ | GC-MS | 0.046 | - | 32.9 | 0.069 | - | 1.2 | -6.5 | 0.65 | - | 0.5 | -29.6 | 0.036 | - | 1.0 |
| Succinic acid | 118.0266 | 10.389 | 17.9 | - | - | Fiehn | $C_4H_6O_4$ | GC-MS | 0.29 | - | -74.8 | 0.56 | - | 1.6 | -84.0 | 0.047 | pBH | 1.8 | -36.5 | 0.12 | - | 1.3 |
| **Fatty Acyls [FA]** | | | | | | | | | | | | | | | | | | | | | | |
| *Carnitines* | | | | | | | | | | | | | | | | | | | | | | |
| Acetyl-L-carnitine | 203.1160 | 0.74 | 1.4 | [M+H]+ | 1 | DB | $C_9H_{17}NO_4$ | LC-MS (+) | $2.0X10^{-5}$ | pBH | -23.9 | 0.021 | pBH | 1.5 | -39.7 | $2.3X10^{-4}$ | pBH | 3.1 | -20.9 | 0.0014 | - | 1.3 |
| Tetradecanoylcarnitine | 371.3063 | 1.71 | 3.4 | [M+H]+ | 7 | MS/MS | $C_{21}H_{41}NO_4$ | LC-MS (+) | 0.012 | - | 36.9 | 0.013 | - | 1.5 | 5.5 | 0.57 | - | 0.8 | -22.9 | 0.033 | - | 1.3 |
| O-arachidonoylcarnitine | 447.3355 | 2.21 | 4.7 | [M+H]+ | 1 | DB | $C_{27}H_{45}NO_4$ | LC-MS (+) | $8.0X10^{-4}$ | pBH | -38.3 | 0.0012 | - | 2.2 | -75.3 | $3.4X10^{-8}$ | pBH | 5.4 | -60.0 | 0.015 | pBH | 2.5 |
| Palmitoylcarnitine | 399.3374 | 2.41 | 0.8 | [M+H]+ | 6 | MS/MS | $C_{23}H_{45}NO_4$ | LC-MS (+) | 0.12 | - | 37.2 | 0.039 | - | 1.4 | 11.8 | 0.49 | - | 1.0 | -18.5 | 0.069 | - | 1.2 |
| Stearoylcarnitine | 427.3668 | 3.49 | 1.3 | [M+H]+ | 1 | MS/MS | $C_{25}H_{49}NO_4$ | LC-MS (+) | 0.055 | - | 32.1 | 0.069 | - | 1.4 | 2.9 | 0.81 | - | 0.8 | -22.1 | 0.035 | - | 1.3 |
| *Eicosanoids* | | | | | | | | | | | | | | | | | | | | | | |
| LTF4 | 585.3029 | 2.66 | 4.9 | [M+NH4]+ | 10 | DB | $C_{28}H_{44}N_2O_8S$ | LC-MS (+) | 0.11 | - | 22.9 | 0.041 | - | 1.2 | 6.9 | 0.52 | - | 0.9 | -13.0 | 0.16 | - | 1.0 |
| *Fatty acids and conjugates* | | | | | | | | | | | | | | | | | | | | | | |
| DHA | 328.2423 | 4.01 | 4.0 | [M-H]- | 7 | MS/MS | $C_{22}H_{32}O_2$ | LC-MS (-) | 0.0010 | - | 5.6 | 0.61 | - | 0.9 | -30.8 | 0.0013 | - | 2.9 | -34.5 | $4.8X10^{-4}$ | - | 1.9 |
| Arachidonic acid | 304.2418 | 4.10 | 4.0 | [M-H]- | 5 | MS/MS | $C_{20}H_{32}O_2$ | LC-MS (-) | 0.0023 | - | -6.0 | 0.64 | - | 0.8 | -40.1 | 0.0018 | - | 3.3 | -36.2 | 0.0026 | - | 1.9 |
| 5-Methyltetrahydrofolic acid | 495.1699 | 4.10 | 5.4 | [M+Cl]- | 15 | DB | $C_{20}H_{25}N_7O_6$ | LC-MS (-) | 0.0017 | - | -2.2 | 0.86 | - | 0.8 | -38.0 | 0.0022 | - | 3.1 | -36.5 | $9.6X10^{-4}$ | - | 2.0 |
| n-Hydroxy stearic acid | 300.2676 | 4.59 | 4.7 | [M-H]- | 4 | DB | $C_{18}H_{36}O_3$ | LC-MS (-) | $1.4X10^{-7}$ | pBH | 670.7 | $7.9X10^{-4}$ | pBH | 4.0 | 935.5 | $4.7X10^{-7}$ | - | 8.2 | 34.4 | 0.029 | - | 1.4 |
| Stearic acid | 284.2726 | 6.58 | 5.2 | [M-H]- | 4 | DB | $C_{18}H_{36}O_2$ | LC-MS (-) | $4.0X10^{-4}$ | - | -1.8 | 0.86 | - | 0.8 | -36.3 | 0.0012 | - | 3.2 | -35.1 | 0.045 | pBH | 2.1 |
| *Fatty amides* | | | | | | | | | | | | | | | | | | | | | | |
| Oleamide | 281.2719 | 3.65 | 4.8 | [M+H]+ | 0 | MS/MS | $C_{18}H_{35}NO$ | LC-MS (+) | 0.21 | - | 117.6 | 0.038 | - | 1.9 | 88.1 | 0.036 | - | 2.3 | -13.5 | 0.87 | - | 0.4 |
| *Fatty esters* | | | | | | | | | | | | | | | | | | | | | | |
| Palmitic acid methyl ester | 270.2559 | 17.635 | 7.1 | - | - | NIST | $C_{17}H_{34}O_2$ | GC-MS | 0.040 | pBH | 21.9 | 0.061 | - | 1.1 | -15.7 | 0.10 | - | 0.9 | -30.8 | 0.016 | pBH | 1.3 |
| **Gamma-keto acids and derivatives** | | | | | | | | | | | | | | | | | | | | | | |
| α-Ketoglutaric acid | 146.0215 | 13.984 | 7.2 | - | - | Fiehn | $C_5H_6O_5$ | GC-MS | 0.0014 | pBH | 26.0 | 0.0026 | pBH | 1.2 | 2.3 | 0.94 | - | 0.6 | -19.8 | 0.0062 | pBH | 1.2 |
| **Glycerolipids [GL]** | | | | | | | | | | | | | | | | | | | | | | |
| *Monoradylglycerols* | | | | | | | | | | | | | | | | | | | | | | |
| MG(22.6) | 440.2340 | 3.11 | 2.5 | [M+K]+ | 3 | DB | $C_{25}H_{38}O_4$ | LC-MS (+) | 0.016 | - | 36.6 | 0.014 | - | 1.4 | 9.2 | 0.30 | - | 0.9 | -20.0 | 0.051 | - | 1.1 |
| MG(16:0) | 330.2787 | 3.58 | 2.1 | [M+H]+ | 5 | DB | $C_{19}H_{38}O_4$ | LC-MS (+) | 0.0064 | - | 30.9 | 0.010 | - | 1.5 | 0.7 | 0.94 | - | 0.8 | -23.1 | 0.0076 | - | 1.3 |
| MG(18:1/0:0/0:0) | 356.2944 | 4.00 | 1.3 | [M+H]+ | 5 | MS/MS | $C_{21}H_{40}O_4$ | LC-MS (+) | 0.014 | - | 21.8 | 0.089 | - | 1.2 | -11.4 | 0.22 | - | 1.0 | -27.3 | 0.0070 | - | 1.4 |
| MG(18:0/0:0/0:0) | 358.3100 | 5.13 | 3.2 | [M+H]+ | 5 | MS/MS | $C_{21}H_{42}O_4$ | LC-MS (+) | 0.11 | - | 27.5 | 0.042 | - | 1.3 | 18.4 | 0.12 | - | 1.2 | -7.1 | 0.52 | - | 0.9 |
| MG(20:1) | 406.3078 | 5.56 | 6.9 | [M+Na]+ | 5 | DB | $C_{23}H_{44}O_4$ | LC-MS (+) | 0.069 | - | 15.4 | 0.26 | - | 1.0 | -16.1 | 0.22 | - | 1.1 | -27.3 | 0.014 | - | 1.4 |

| Compound | Mass | RT | CV | Adduct | Error | ID source | Formula | Analytical platform | p value ANOVA | pBH | % change | p value t test | p BH | VIP | % change | p value t test | p BH | VIP | % change | p value t test | p BH | VIP |
|---|---|---|---|---|---|---|---|---|---|---|---|---|---|---|---|---|---|---|---|---|---|---|
| | | | | | | | | | | | **2.5h *vs* 35 min** | | | | **5.5h *vs* 35 min** | | | | **5.5h *vs* 2.5h** | | | |
| *Diradylglycerols* | | | | | | | | | | | | | | | | | | | | | | |
| DG(36:4) | 638.4871 | 11.46 | 4.2 | [M+Na]+ | 3 | DB | $C_{39}H_{68}O_5$ | LC-MS (+) | 0.0028 | - | 31.5 | 0.0024 | - | 1.5 | 18.3 | 0.012 | - | 1.5 | -10.1 | 0.15 | - | 0.8 |
| DG(18:1/20:4) | 624.5122 | 12.00 | 2.8 | [M+H-H2O]+ | 0 | MS/MS | $C_{41}H_{70}O_5$ | LC-MS (+) | 0.023 | - | 30.4 | 0.011 | - | 1.5 | 10.9 | 0.30 | - | 1.1 | -14.9 | 0.076 | - | 1.0 |
| DG(38:5) | 664.5036 | 12.00 | 3.2 | [M+Na]+ | 1 | DB | $C_{41}H_{70}O_5$ | LC-MS (+) | 0.011 | - | 26.9 | 0.0032 | - | 1.4 | 14.7 | 0.082 | - | 1.2 | -9.6 | 0.17 | - | 0.8 |
| DG(34:1) | 576.5129 | 12.38 | 4.1 | [M+H-H2O]+ | 2 | DB | $C_{37}H_{70}O_5$ | LC-MS (+) | 0.058 | - | 19.4 | 0.11 | - | 1.1 | -4.7 | 0.57 | - | 0.7 | -20.2 | 0.023 | - | 1.2 |
| DG(40:6) | 650.5287 | 13.05 | 5.1 | [M+H-H2O]+ | 2 | DB | $C_{43}H_{72}O_5$ | LC-MS (+) | 0.020 | - | 40.5 | 0.020 | - | 1.6 | 10.9 | 0.34 | - | 1.1 | -21.1 | 0.054 | - | 1.2 |
| 1-Hexadecyl-2-arachidonoyl-glycerol | 602.5291 | 13.18 | 3.4 | [M+H]+ | 3 | DB | $C_{39}H_{70}O_4$ | LC-MS (+) | 0.092 | - | 21.4 | 0.15 | - | 1.3 | -7.2 | 0.58 | - | 1.0 | -23.6 | 0.035 | - | 1.3 |
| DG(38:4) | 626.5286 | 13.47 | 9.8 | [M+H-H2O]+ | 2 | DB | $C_{41}H_{72}O_5$ | LC-MS (+) | 0.040 | - | 24.1 | 0.11 | - | 1.2 | -7.3 | 0.46 | - | 0.9 | -25.3 | 0.017 | - | 1.3 |
| DG(38:6) | 622.4965 | 13.68 | 5.2 | [M+H-H2O]+ | 0 | DB | $C_{41}H_{68}O_5$ | LC-MS (+) | 0.027 | - | 22.2 | 0.10 | - | 1.3 | -13.0 | 0.31 | - | 1.3 | -28.8 | 0.010 | - | 1.6 |
| DG(36:4) | 598.4959 | 13.83 | 7.3 | [M+H-H2O]+ | 1 | DB | $C_{39}H_{68}O_5$ | LC-MS (+) | 0.040 | - | 24.0 | 0.080 | - | 1.3 | -5.6 | 0.63 | - | 0.9 | -23.8 | 0.016 | - | 1.3 |
| DG(35:0) | 648.5132 | 14.21 | 10.0 | [M+K]+ | 6 | DB | $C_{38}H_{74}O_5$ | LC-MS (+) | 0.014 | - | 18.8 | 0.076 | - | 1.2 | -10.8 | 0.29 | - | 1.1 | -24.9 | 0.0035 | - | 1.4 |
| DG(40:4) | 654.5573 | 14.58 | 4.7 | [M+H-H2O]+ | 2 | DB | $C_{43}H_{76}O_5$ | LC-MS (+) | 0.10 | - | 17.0 | 0.18 | - | 1.2 | -6.7 | 0.56 | - | 0.9 | -20.3 | 0.025 | - | 1.2 |
| DG(34:8) | 705.5336 | 15.42 | 4.0 | [M+NH4]+ | 0 | DB | $C_{45}H_{68}O_5$ | LC-MS (+) | 0.0067 | pBH | 41.7 | 0.017 | pBH | 1.7 | 18.1 | 0.032 | - | 1.5 | -16.6 | 0.010 | - | 1.0 |
| DG(38:4) | 626.5291 | 15.56 | 5.6 | [M+H-H2O]+ | 2 | DB | $C_{41}H_{72}O_5$ | LC-MS (+) | 0.0476 | - | 19.3 | 0.18 | - | 1.2 | -13.0 | 0.31 | - | 1.2 | -27.1 | 0.012 | - | 1.5 |
| DG(40:8) | 681.5340 | 15.60 | 4.6 | [M+NH4]+ | 1 | DB | $C_{43}H_{68}O_5$ | LC-MS (+) | 0.0418 | - | 1.6 | 0.88 | - | 0.7 | -19.8 | 0.051 | - | 1.6 | -21.0 | 0.014 | - | 1.2 |
| DG(36:3) | 640.5069 | 16.01 | 3.7 | [M+Na]+ | 4 | DB | $C_{39}H_{70}O_5$ | LC-MS (+) | 0.0043 | - | 17.4 | 0.013 | - | 1.0 | 21.0 | 0.0026 | - | 1.7 | 3.1 | 0.55 | - | 0.5 |
| DG(34:0) | 604.5431 | 16.48 | 5.1 | [M+Na]+ | 4 | DB | $C_{37}H_{74}O_4$ | LC-MS (+) | 0.0469 | - | 18.5 | 0.17 | - | 1.2 | -11.2 | 0.31 | - | 1.1 | -25.1 | 0.015 | - | 1.4 |
| DG(40:7) | 688.5059 | 16.55 | 4.3 | [M+Na]+ | 2 | DB | $C_{43}H_{70}O_5$ | LC-MS (+) | $1.1X10^{-4}$ | pBH | 22.6 | 0.019 | pBH | 1.3 | 36.8 | 0.0012 | pBH | 2.4 | 11.6 | 0.0076 | - | 0.8 |
| DG(32:0) | 585.5337 | 16.67 | 4.2 | [M+NH4]+ | 1 | DB | $C_{35}H_{68}O_5$ | LC-MS (+) | 0.0245 | - | 31.1 | 0.016 | - | 1.4 | 20.2 | 0.035 | - | 1.5 | -8.3 | 0.34 | - | 0.8 |
| DG(40:4) | 654.5576 | 16.70 | 5.0 | [M+H-H2O]+ | 2 | DB | $C_{43}H_{76}O_5$ | LC-MS (+) | 0.0513 | - | 22.6 | 0.14 | - | 1.3 | -10.6 | 0.423 | - | 1.1 | -27.1 | 0.015 | - | 1.5 |
| DG(32:0) | 590.4901 | 16.72 | 2.6 | [M+Na]+ | 2 | DB | $C_{35}H_{68}O_5$ | LC-MS (+) | $1.1X10^{-4}$ | pBH | 16.3 | 0.013 | pBH | 1.2 | 22.7 | $2.0X10^{-5}$ | pBH | 2.0 | 5.5 | 0.090 | - | 0.6 |
| DG(38:5) | 664.5016 | 16.95 | 3.4 | [M+Na]+ | 4 | DB | $C_{41}H_{70}O_5$ | LC-MS (+) | 0.0015 | pBH | 22.8 | 0.041 | pBH | 1.2 | 35.6 | 0.0048 | pBH | 2.4 | 10.4 | 0.069 | - | 0.8 |
| DG(38:3) | 668.5389 | 17.60 | 3.5 | [M+Na]+ | 5 | DB | $C_{41}H_{74}O_5$ | LC-MS (+) | 0.032 | - | 27.7 | 0.024 | - | 1.3 | 25.3 | 0.024 | - | 1.7 | -1.9 | 0.82 | - | 0.7 |
| DG(40:6) | 706.4951 | 17.63 | 2.9 | [M+K]+ | 2 | DB | $C_{43}H_{72}O_5$ | LC-MS (+) | 0.0019 | - | 6.3 | 0.19 | - | 0.5 | 29.9 | 0.023 | pBH | 2.0 | 22.2 | 0.010 | - | 1.1 |
| DG(34:0) | 618.5212 | 18.22 | 3.9 | [M+Na]+ | 2 | DB | $C_{37}H_{72}O_5$ | LC-MS (+) | 0.0011 | pBH | 33.3 | 0.011 | pBH | 1.6 | 38.3 | $8.5X10^{-6}$ | pBH | 2.5 | 3.7 | 0.078 | - | 0.6 |
| DG(38:0) | 668.5366 | 18.37 | 3.2 | [M+Na]+ | 1 | DB | $C_{41}H_{74}O_5$ | LC-MS (+) | 0.32 | - | 28.9 | 0.13 | - | 1.1 | 14.2 | 0.017 | - | 1.3 | -11.4 | 0.54 | - | 0.9 |
| *Triradylglycerols* | | | | | | | | | | | | | | | | | | | | | | |
| TG(37:0) | 669.5818 | 18.80 | 2.7 | [M+NH4]+ | 14 | DB | $C_{40}H_{76}O_6$ | LC-MS (+) | $1.0X10^{-5}$ | pBH | 99.1 | 0.0013 | pBH | 2.7 | 122.7 | 0.007 | - | 4.0 | 11.9 | 0.19 | - | 0.9 |
| TG(16:0/22:6/22:6) | 967.7624 | 24.22 | 3.5 | [M+NH4]+ | 1 | MS/MS | $C_{45}H_{68}O_5$ | LC-MS (+) | 0.022 | pBH | 41.4 | 0.0011 | - | 1.7 | 44.6 | 0.025 | - | 2.4 | 2.3 | 0.78 | - | 0.7 |
| TG(58:10) | 948.7197 | 24.40 | 2.2 | [M+Na]+ | 1 | DB | $C_{61}H_{98}O_6$ | LC-MS (+) | 0.0091 | - | 37.1 | 0.011 | - | 1.5 | 6.5 | 0.878 | - | 0.9 | -22.4 | 0.0086 | - | 1.3 |
| TG(56:8) | 924.7209 | 24.60 | 1.5 | [M+Na]+ | 3 | DB | $C_{59}H_{98}O_6$ | LC-MS (+) | 0.011 | - | 7.0 | 0.60 | - | 0.9 | -26.8 | 0.021 | - | 1.9 | -31.6 | 0.0033 | - | 1.6 |
| TG(42:8) | 1000.7521 | 25.17 | 5.8 | [M+Na]+ | 2 | DB | $C_{65}H_{102}O_6$ | LC-MS (+) | $6.7X10^{-4}$ | pBH | 58.0 | 0.0058 | pBH | 2.1 | 56.3 | 0.004 | pBH | 2.8 | -1.0 | 0.89 | - | 0.6 |
| TG(54:5) | 918.7078 | 25.34 | 5.7 | [M+K]+ | 0 | DB | $C_{57}H_{100}O_6$ | LC-MS (+) | 0.0061 | - | 5.3 | 0.58 | - | 0.7 | -21.8 | 0.011 | - | 1.8 | -25.7 | 0.0025 | - | 1.4 |
| TG(54:5) | 902.7342 | 25.35 | 3.4 | [M+Na]+ | 0 | DB | $C_{57}H_{100}O_6$ | LC-MS (+) | 0.037 | - | 5.1 | 0.62 | - | 0.8 | -17.3 | 0.067 | - | 1.4 | -21.3 | 0.0079 | - | 1.3 |
| TG(60:10) | 992.7264 | 25.39 | 4.8 | [M+K]+ | 3 | DB | $C_{63}H_{102}O_6$ | LC-MS (+) | 0.013 | - | 21.4 | 0.030 | - | 1.2 | -4.9 | 0.58 | - | 0.8 | -21.7 | 0.0071 | - | 1.3 |
| TG(22:6/16:0/18:0) | 923.7945 | 25.85 | 2.7 | [M+NH4]+ | 0 | MS/MS | $C_{59}H_{102}O_6$ | LC-MS (+) | 0.059 | - | 20.1 | 0.022 | - | 1.2 | 14.7 | 0.11 | - | 1.2 | -4.5 | 0.49 | - | 0.7 |
| TG(58:7) | 949.8081 | 26.23 | 2.5 | [M+NH4]+ | 2 | DB | $C_{61}H_{104}O_6$ | LC-MS (+) | 0.060 | - | 20.3 | 0.026 | - | 1.2 | 9.0 | 0.29 | - | 1.0 | -9.3 | 0.15 | - | 0.8 |
| TG(56:5) | 930.7664 | 26.54 | 3.7 | [M+Na]+ | 1 | DB | $C_{61}H_{102}O_6$ | LC-MS (+) | 0.022 | - | 9.2 | 0.30 | - | 0.8 | -13.8 | 0.11 | - | 1.3 | -21.1 | 0.0037 | - | 1.3 |
| **Glycerophospholipids [GP]** | | | | | | | | | | | | | | | | | | | | | | |
| *Glycerophosphates* | | | | | | | | | | | | | | | | | | | | | | |
| LysoPA(22:4) | 486.2838 | 9.78 | 7.2 | [M-H]- | 19 | DB | $C_{25}H_{43}O_7P$ | LC-MS (-) | 0.0095 | - | 25.0 | 0.015 | - | 1.5 | -1.7 | 0.82 | - | 0.8 | -21.4 | 0.0099 | - | 1.4 |
| PA(43:6) | 790.5500 | 14.28 | 3.2 | [M+H]+ | 2 | DB | $C_{46}H_{79}O_8P$ | LC-MS (+) | 0.044 | - | 18.1 | 0.10 | - | 1.2 | -6.2 | 0.51 | - | 0.9 | -20.6 | 0.016 | - | 1.2 |
| PA(40:2) | 773.6009 | 15.38 | 7.6 | [M+NH4]+ | 10 | DB | $C_{43}H_{81}O_8P$ | LC-MS (+) | 0.015 | - | 5.1 | 0.67 | - | 0.9 | -24.4 | 0.015 | - | 1.8 | -28.1 | 0.0054 | - | 1.5 |
| PA(34:1) | 674.4899 | 16.40 | 4.0 | [M-H]- | 2 | DB | $C_{37}H_{71}O_8P$ | LC-MS (-) | 0.025 | - | 23.7 | 0.011 | - | 1.3 | 14.1 | 0.10 | - | 1.4 | -7.8 | 0.24 | - | 0.7 |
| *Glycerophosphocholines* | | | | | | | | | | | | | | | | | | | | | | |
| Glycerophosphocholine | 257.1036 | 0.73 | 2.4 | [M+H]+ | 3 | MS/MS | $C_8H_{20}NO_6P$ | LC-MS (+) | $2.6X10^{-7}$ | pBH | -32.3 | 0.006 | pBH | 1.9 | -53.3 | $8.7X10^{-7}$ | pBH | 4.0 | -31.0 | $7.3X10^{-4}$ | - | 1.6 |
| LysoPC(0:0/20:4) | 543.3337 | 2.21 | 2.2 | [M+H]+ | 2 | MS/MS | $C_{28}H_{50}NO_7P$ | LC-MS (+) | 0.077 | - | 25.0 | 0.10 | - | 1.3 | -3.4 | 0.78 | - | 1.0 | -22.7 | 0.048 | - | 1.3 |
| LysoPC(0:0/20:4) | 589.3396 | 2.28 | 3.8 | [M+FA-H]- | 3 | MS/MS | $C_{28}H_{50}NO_7P$ | LC-MS (-) | 0.10 | - | 24.3 | 0.12 | - | 1.4 | -2.8 | 0.81 | - | 1.1 | -21.8 | 0.069 | - | 1.4 |
| LysoPC(0:0/22:6) | 613.3394 | 2.25 | 4.2 | [M+FA-H]- | 3 | MS/MS | $C_{30}H_{50}NO_7P$ | LC-MS (-) | 0.10 | - | 30.6 | 0.069 | - | 1.5 | 5.4 | 0.65 | - | 1.0 | -19.3 | 0.13 | - | 1.3 |
| LysoPC(22:6/0:0) | 613.3400 | 2.41 | 3.7 | [M+FA-H]- | 4 | MS/MS | $C_{30}H_{50}NO_7P$ | LC-MS (-) | 0.13 | - | 33.1 | 0.080 | - | 1.6 | 7.7 | 0.57 | - | 1.2 | -19.1 | 0.18 | - | 1.3 |
| LysoPC(18:2/0:0) | 565.3398 | 2.46 | 5.1 | [M+FA-H]- | 4 | DB | $C_{26}H_{50}NO_7P$ | LC-MS (-) | 0.12 | - | 31.9 | 0.068 | - | 1.6 | 6.6 | 0.64 | - | 1.2 | -19.1 | 0.13 | - | 1.3 |
| LysoPC(20:4/0:0) | 589.3401 | 2.46 | 4.2 | [M+FA-H]- | 4 | MS/MS | $C_{28}H_{50}NO_7P$ | LC-MS (-) | 0.16 | - | 30.3 | 0.12 | - | 1.5 | 2.6 | 0.85 | - | 1.2 | -21.3 | 0.14 | - | 1.4 |
| LysoPC(0:0/16:0) | 541.3405 | 2.51 | 4.0 | [M+FA-H]- | 5 | MS/MS | $C_{24}H_{50}NO_7P$ | LC-MS (-) | 0.087 | - | 32.5 | 0.057 | - | 1.5 | 7.2 | 0.56 | - | 1.1 | -19.1 | 0.13 | - | 1.3 |
| LysoPC(0:0/18:1) | 567.3554 | 2.85 | 3.7 | [M+FA-H]- | 4 | MS/MS | $C_{26}H_{52}NO_7P$ | LC-MS (-) | 0.11 | - | 30.1 | 0.072 | - | 1.5 | 6.6 | 0.57 | - | 1.1 | -18.1 | 0.15 | - | 1.2 |
| LysoPC(0:0/18:0) | 569.3714 | 3.66 | 3.6 | [M-H]- | 4 | MS/MS | $C_{26}H_{54}NO_7P$ | LC-MS (-) | 0.086 | - | 33.6 | 0.039 | - | 1.6 | 14.8 | 0.24 | - | 1.5 | -14.1 | 0.27 | - | 1.1 |
| LysoPC(0:0/18:0) | 523.3652 | 3.53 | 3.6 | [M+H]+ | 3 | DB | $C_{26}H_{54}NO_7P$ | LC-MS (+) | 0.10 | - | 34.2 | 0.045 | - | 1.5 | 15.0 | 0.27 | - | 1.2 | -14.3 | 0.25 | - | 1.1 |
| LysoPC(18:0/0:0) | 569.3712 | 4.00 | 3.6 | [M+FA-H]- | 4 | MS/MS | $C_{26}H_{54}NO_7P$ | LC-MS (-) | 0.056 | - | 37.3 | 0.027 | - | 1.7 | 14.9 | 0.24 | - | 1.5 | -16.3 | 0.20 | - | 1.2 |
| LysoPC(18:0/0:0) | 523.3648 | 3.86 | 1.6 | [M+H]+ | 2 | MS/MS | $C_{26}H_{54}NO_7P$ | LC-MS (+) | 0.082 | - | 39.6 | 0.046 | - | 1.6 | 13.0 | 0.38 | - | 1.2 | -19.1 | 0.16 | - | 1.2 |
| LysoPC(20:3/0:0) | 525.2819 | 3.89 | 3.7 | [M+Na]+ | 3 | DB | $C_{25}H_{46}NO_7P$ | LC-MS (+) | 0.052 | - | 30.5 | 0.023 | - | 1.4 | 7.4 | 0.54 | - | 0.9 | -17.7 | 0.092 | - | 1.2 |

| Compound | Mass | RT | CV | Adduct | Error | ID source | Formula | Analytical platform | p value ANOVA | pBH | % change | p value t test | p BH | VIP | % change | p value t test | p BH | VIP | % change | p value t test | p BH | VIP |
|---|---|---|---|---|---|---|---|---|---|---|---|---|---|---|---|---|---|---|---|---|---|---|
| | | | | | | | | | | | 2.5h *vs* 35 min | | | | 5.5h *vs* 35 min | | | | 5.5h *vs* 2.5h | | | |
| PC(36:7) | 792.5379 | 12.32 | 2.5 | [M+NH4) | 5 | DB | $C_{44}H_{74}NO_8P$ | LC-MS (+) | 0.022 | - | 21.9 | 0.090 | - | 1.3 | -9.3 | 0.41 | - | 1.0 | -25.6 | 0.0038 | - | 1.4 |
| PC(22:6/22:6) | 877.5622 | 12.73 | 2.4 | [M+H]+ | 0 | MS/MS | $C_{52}H_{80}NO_8P$ | LC-MS (+) | 0.033 | - | 29.3 | 0.029 | - | 1.4 | 5.2 | 0.57 | - | 0.8 | -18.6 | 0.058 | - | 1.1 |
| PC(20:4/22:6) | 853.5621 | 12.90 | 2.5 | [M+H]+ | 0 | MS/MS | $C_{50}H_{80}NO_8P$ | LC-MS (+) | 0.022 | - | 23.5 | 0.026 | - | 1.3 | 0.2 | 0.98 | - | 0.8 | -18.9 | 0.019 | - | 1.2 |
| PC(35:3) | 805.5383 | 12.95 | 3.4 | [M+Cl]- | 1 | DB | $C_{43}H_{80}NO_8P$ | LC-MS (-) | 0.061 | - | 23.4 | 0.075 | - | 1.4 | -0.6 | 0.95 | - | 0.8 | -19.5 | 0.047 | - | 1.3 |
| PC(16:0/15:0) | 719.5546 | 13.02 | 4.7 | [M+H]+ | 11 | MS/MS | $C_{39}H_{78}NO_8P$ | LC-MS (+) | 0.066 | - | 22.2 | 0.094 | - | 1.1 | -1.5 | 0.86 | - | 0.6 | -19.5 | 0.049 | - | 1.1 |
| PC(16:1/16:0) | 731.5475 | 13.06 | 2.1 | [M+H]+ | 1 | MS/MS | $C_{40}H_{78}NO_8P$ | LC-MS (+) | 0.065 | - | 17.6 | 0.12 | - | 1.1 | -6.3 | 0.53 | - | 0.9 | -20.3 | 0.027 | - | 1.2 |
| PC(18:2/22:6) | 829.5627 | 13.06 | 3.6 | [M+H]+ | 1 | MS/MS | $C_{48}H_{80}NO_8P$ | LC-MS (+) | 0.029 | - | 21.6 | 0.056 | - | 1.3 | -3.7 | 0.70 | - | 0.8 | -20.8 | 0.012 | - | 1.3 |
| PC(44:12) | 923.5666 | 13.07 | 4.9 | [M+FA-H]- | 1 | MS/MS | $C_{52}H_{80}NO_8P$ | LC-MS (-) | 0.036 | - | 26.2 | 0.045 | - | 1.4 | 5.4 | 0.53 | - | 0.8 | -16.5 | 0.13 | - | 1.2 |
| PC(22:6/16:0) | 805.5630 | 13.15 | 1.2 | [M+H]+ | 1 | MS/MS | $C_{46}H_{80}NO_8P$ | LC-MS (+) | 0.023 | - | 12.5 | 0.30 | - | 1.1 | -17.6 | 0.14 | - | 1.3 | -26.8 | 0.0022 | - | 1.5 |
| PC(20:4/22:6) / PC(22:6/20:4) | 899.5679 | 13.23 | 5.9 | [M+FA-H]- | 0 | MS/MS | $C_{50}H_{80}NO_8P$ | LC-MS (-) | 0.0060 | - | 24.9 | 0.013 | - | 1.4 | -2.3 | 0.79 | - | 0.9 | -21.8 | 0.0050 | - | 1.5 |
| PC(48:6) | 851.5685 | 13.47 | 3.8 | [M+FA-H]- | 1 | DB | $C_{46}H_{80}NO_8P$ | LC-MS (-) | 0.022 | - | 11.3 | 0.28 | - | 1.0 | -15.0 | 0.13 | - | 1.5 | -23.7 | 0.0032 | - | 1.5 |
| PC(16:0/16:1) | 731.5477 | 13.51 | 8.4 | [M+H]+ | 2 | MS/MS | $C_{40}H_{78}NO_8P$ | LC-MS (+) | 0.0020 | - | 21.2 | 0.021 | - | 1.2 | -9.6 | 0.23 | - | 1.1 | -25.4 | 0.0012 | - | 1.4 |
| PC(44:11) | 879.5751 | 13.75 | 4.0 | [M+H]+ | 3 | DB | $C_{52}H_{82}NO_8P$ | LC-MS (+) | 0.0092 | - | 21.4 | 0.0063 | - | 1.3 | 4.2 | 0.56 | - | 0.7 | -14.2 | 0.016 | - | 1.0 |
| PC(22:6/18:1) | 831.5778 | 13.78 | 5.8 | [M+H]+ | 0 | MS/MS | $C_{48}H_{82}NO_8P$ | LC-MS (+) | 0.042 | - | 16.6 | 0.18 | - | 1.1 | -11.5 | 0.28 | - | 1.1 | -24.1 | 0.013 | - | 1.3 |
| PC(18:1/18:2) | 783.5788 | 14.31 | 5.1 | [M+H]+ | 1 | MS/MS | $C_{44}H_{82}NO_8P$ | LC-MS (+) | 0.010 | - | 19.6 | 0.033 | - | 1.2 | -7.6 | 0.41 | - | 1.0 | -22.8 | 0.0035 | - | 1.4 |
| PC(20:4/18:1) | 853.5828 | 14.36 | 5.2 | [M+FA-H]- | 1 | MS/MS | $C_{46}H_{82}NO_8P$ | LC-MS (-) | 0.00029 | - | 13.0 | 0.097 | - | 1.1 | -19.9 | 0.017 | - | 2.0 | -29.1 | 0.045 | pBH | 1.8 |
| PC(35:3) | 769.5635 | 16.05 | 6.4 | [M+H]+ | 2 | DB | $C_{43}H_{80}NO_8P$ | LC-MS (+) | 0.027 | - | 16.9 | 0.22 | - | 1.1 | -16.5 | 0.13 | - | 1.3 | -28.6 | 0.0086 | - | 1.5 |
| PC(40:3) | 861.6252 | 16.90 | 4.9 | [M+Na]+ | 3 | DB | $C_{48}H_{90}NO_8P$ | LC-MS (+) | 0.071 | - | 23.4 | 0.077 | - | 1.3 | -2.5 | 0.83 | - | 0.9 | -20.9 | 0.039 | - | 1.3 |
| PC(40:4) | 837.6232 | 17.09 | 6.7 | [M+H]+ | 2 | DB | $C_{48}H_{88}NO_8P$ | LC-MS (+) | 0.039 | - | 16.7 | 0.17 | - | 1.0 | -9.9 | 0.27 | - | 0.9 | -22.8 | 0.013 | - | 1.2 |
| PC(18:0/18:0) | 789.6259 | 17.42 | 3.8 | [M+H]+ | 1 | MS/MS | $C_{44}H_{88}NO_8P$ | LC-MS (+) | 0.082 | - | 18.2 | 0.16 | - | 1.1 | -6.2 | 0.53 | - | 0.9 | -20.6 | 0.035 | - | 1.2 |
| PC(18:0/20:1) | 815.6417 | 17.75 | 2.6 | [M+H]+ | 1 | MS/MS | $C_{46}H_{90}NO_8P$ | LC-MS (+) | 0.080 | - | 19.4 | 0.13 | - | 1.2 | -6.3 | 0.54 | - | 0.9 | -21.6 | 0.025 | - | 1.2 |
| PC(42:4) | 865.6502 | 18.50 | 4.2 | [M+H]+ | 7 | DB | $C_{50}H_{92}NO_8P$ | LC-MS (+) | 0.046 | pBH | 27.7 | 0.010 | - | 1.4 | -4.4 | 0.56 | - | 0.8 | -25.1 | 0.0012 | - | 1.4 |
| PC(40:2) | 877.6342 | 18.96 | 4.8 | [M+Cl]- | 2 | DB | $C_{48}H_{92}NO_8P$ | LC-MS (-) | 0.020 | - | 24.4 | 0.021 | - | 1.3 | 2.1 | 0.75 | - | 0.8 | -17.9 | 0.069 | - | 1.2 |
| PC(18:1/24:0) | 917.7086 | 20.96 | 4.0 | [M+FA-H]- | 0 | MS/MS | $C_{50}H_{98}NO_8P$ | LC-MS (-) | 0.061 | - | 24.1 | 0.091 | - | 1.3 | -2.1 | 0.81 | - | 0.8 | -21.2 | 0.048 | - | 1.3 |
| PC(O-17:0/17:0) | 747.6150 | 16.92 | 4.2 | [M+H]+ | 1 | MS/MS | $C_{42}H_{86}NO_7P$ | LC-MS (+) | 0.069 | - | 14.9 | 0.21 | - | 1.1 | -9.5 | 0.35 | - | 1.0 | -21.3 | 0.018 | - | 1.3 |
| PC(P-38:1) / PC(O-38:2) | 799.6532 | 14.76 | 2.5 | [M+H]+ | 10 | DB | $C_{46}H_{90}NO_7P$ | LC-MS (+) | 0.017 | - | 34.3 | 0.020 | - | 1.6 | -1.2 | 0.93 | - | 1.0 | -26.4 | 0.011 | - | 1.5 |
| PC(P-35:3) / PC(O-35:4) | 753.5666 | 16.18 | 5.2 | [M+H]+ | 1 | DB | $C_{43}H_{80}NO_7P$ | LC-MS (+) | 0.11 | - | 11.4 | 0.35 | - | 1.0 | -13.9 | 0.28 | - | 1.3 | -22.7 | 0.028 | - | 1.4 |
| PC(P-32:2) / PC(O-32:3) | 899.6985 | 17.84 | 9.1 | [M+FA-H]- | 1 | DB | $C_{50}H_{96}NO_7P$ | LC-MS (-) | 0.060 | - | 23.9 | 0.076 | - | 1.4 | -2.7 | 0.79 | - | 0.9 | -21.5 | 0.045 | - | 1.3 |
| PC(P-44:5) / PC(O-44:6) | 943.6694 | 19.05 | 5.0 | [M-H+HCOONa]- | 6 | DB | $C_{52}H_{94}NO_7P$ | LC-MS (-) | 0.089 | - | 20.8 | 0.14 | - | 1.3 | -3.9 | 0.70 | - | 0.8 | -20.5 | 0.041 | - | 1.2 |
| PC(20:1) / PC(P-20:0) / PC(O-20:1) | 595.3867 | 4.37 | 7.6 | [M+FA-H]- | 3 | DB | $C_{28}H_{56}NO_7P$ | LC-MS (-) | 0.15 | - | 31.5 | 0.10 | - | 1.5 | 6.4 | 0.61 | - | 1.1 | -19.1 | 0.20 | - | 1.2 |
| *Glycerophosphoethanolamines* | | | | | | | | | | | | | | | | | | | | | | |
| sn-glycero-3-Phosphoethanolamine | 215.0574 | 0.73 | 4.7 | [M-H]- | 7 | DB | $C_5H_{14}NO_6P$ | LC-MS (-) | 0.0015 | - | 0.8 | 0.87 | - | 0.7 | -24.1 | 0.0032 | - | 2.2 | -24.7 | $8.6X10^{-4}$ | - | 1.6 |
| LysoPE(0:0/22:6) | 525.2881 | 2.28 | 3.7 | [M-H]- | 5 | MS/MS | $C_{27}H_{44}NO_7P$ | LC-MS (-) | 0.094 | - | 30.1 | 0.069 | - | 1.5 | 3.8 | 0.75 | - | 1.0 | -20.2 | 0.11 | - | 1.3 |
| LysoPE(22:6/0:0) | 525.2897 | 2.44 | 4.2 | [M-H]- | 8 | DB | $C_{27}H_{44}NO_7P$ | LC-MS (-) | 0.12 | - | 32.3 | 0.066 | - | 1.6 | 7.6 | 0.59 | - | 1.2 | -18.6 | 0.16 | - | 1.3 |
| LysoPE(0:0/20:4) | 501.2879 | 2.31 | 4.2 | [M-H]- | 0 | MS/MS | $C_{25}H_{44}NO_7P$ | LC-MS (-) | 0.082 | - | 29.6 | 0.071 | - | 1.5 | 2.3 | 0.85 | - | 1.1 | -21.1 | 0.081 | - | 1.4 |
| LysoPE(20:4/0:0) | 501.2880 | 2.48 | 3.9 | [M-H]- | 5 | DB | $C_{25}H_{44}NO_7P$ | LC-MS (-) | 0.13 | - | 33.2 | 0.083 | - | 1.6 | 6.5 | 0.65 | - | 1.2 | -20.0 | 0.15 | - | 1.4 |
| LysoPE(0:0/16:0) | 453.2879 | 2.55 | 4.1 | [M-H]- | 5 | MS/MS | $C_{21}H_{44}NO_7P$ | LC-MS (-) | 0.086 | - | 35.2 | 0.046 | - | 1.6 | 12.0 | 0.36 | - | 1.3 | -17.2 | 0.18 | - | 1.2 |
| LysoPE(16:0/0:0) | 453.2885 | 2.78 | 3.8 | [M-H]- | 7 | MS/MS | $C_{21}H_{44}NO_7P$ | LC-MS (-) | 0.085 | - | 36.7 | 0.045 | - | 1.6 | 13.1 | 0.33 | - | 1.3 | -17.3 | 0.19 | - | 1.3 |
| LysoPE(16:0/0:0) | 453.2871 | 2.69 | 1.8 | [M+H]+ | 3 | MS/MS | $C_{21}H_{44}NO_7P$ | LC-MS (+) | 0.072 | - | 33.4 | 0.030 | - | 1.5 | 12.2 | 0.35 | - | 1.1 | -15.9 | 0.17 | - | 1.1 |
| LysoPE(20:3/0:0) | 503.3026 | 2.86 | 4.2 | [M-H]- | 3 | DB | $C_{25}H_{46}NO_7P$ | LC-MS (-) | 0.13 | - | 36.4 | 0.085 | - | 1.6 | 11.0 | 0.42 | - | 1.3 | -18.6 | 0.37 | - | 1.3 |
| LysoPE(0:0/18:1) | 479.3032 | 2.88 | 4.6 | [M-H]- | 4 | MS/MS | $C_{23}H_{46}NO_7P$ | LC-MS (-) | 0.12 | - | 31.8 | 0.087 | - | 1.5 | 6.8 | 0.56 | - | 1.1 | -19.0 | 0.20 | - | 1.2 |
| LysoPE(18:1/0:0) | 479.3042 | 3.13 | 3.5 | [M-H]- | 6 | MS/MS | $C_{23}H_{46}NO_7P$ | LC-MS (-) | 0.070 | - | 37.1 | 0.065 | - | 1.6 | 10.1 | 0.46 | - | 1.3 | -19.7 | 0.16 | - | 1.3 |
| LysoPE(22:5/0:0) | 527.3033 | 2.99 | 4.8 | [M-H]- | 4 | DB | $C_{27}H_{46}NO_7P$ | LC-MS (-) | 0.13 | - | 28.2 | 0.084 | - | 1.5 | 3.0 | 0.83 | - | 1.1 | -19.7 | 0.12 | - | 1.4 |
| LysoPE(0:0/22:4) | 529.3189 | 3.10 | 3.9 | [M-H]- | 4 | MS/MS | $C_{27}H_{48}NO_7P$ | LC-MS (-) | 0.10 | - | 25.9 | 0.076 | - | 1.4 | 3.3 | 0.76 | - | 1.0 | -18.0 | 0.11 | - | 1.3 |
| LysoPE(22:4/0:0) | 529.3191 | 3.33 | 4.1 | [M-H]- | 4 | MS/MS | $C_{27}H_{48}NO_7P$ | LC-MS (-) | 0.12 | - | 34.6 | 0.072 | - | 1.6 | 8.5 | 0.55 | - | 1.3 | -19.4 | 0.16 | - | 1.3 |
| LysoPE(P-16:0/0:0) | 437.2933 | 3.37 | 3.5 | [M-H]- | 6 | MS/MS | $C_{21}H_{44}NO_6P$ | LC-MS (-) | 0.083 | - | 40.9 | 0.039 | - | 1.7 | 17.2 | 0.25 | - | 1.5 | -16.8 | 0.23 | - | 1.3 |
| LysoPE(0:0/18:0) | 481.3183 | 3.58 | 3.6 | [M+H]+ | 3 | DB | $C_{23}H_{48}NO_7P$ | LC-MS (+) | 0.086 | - | 31.9 | 0.044 | - | 1.4 | 10.3 | 0.42 | - | 1.1 | -16.4 | 0.16 | - | 1.1 |
| LysoPE(0:0/18:0) | 481.3195 | 3.70 | 4.0 | [M-H]- | 6 | MS/MS | $C_{23}H_{48}NO_7P$ | LC-MS (-) | 0.047 | - | 41.2 | 0.047 | - | 1.7 | 14.7 | 0.32 | - | 1.4 | -18.8 | 0.19 | - | 1.3 |
| LysoPE(18:0/0:0) | 481.3191 | 3.89 | 1.7 | [M+H]+ | 5 | MS/MS | $C_{23}H_{48}NO_7P$ | LC-MS (+) | 0.072 | - | 36.1 | 0.037 | - | 1.5 | 10.5 | 0.45 | - | 1.1 | -18.8 | 0.13 | - | 1.2 |
| LysoPE(18:0/0:0) | 481.3212 | 4.03 | 4.3 | [M-H]- | 9 | MS/MS | $C_{23}H_{48}NO_7P$ | LC-MS (-) | 0.055 | - | 35.4 | 0.047 | - | 1.6 | 12.7 | 0.33 | - | 1.3 | -16.8 | 0.19 | - | 1.2 |
| LysoPE(20:1/0:0) | 507.3347 | 4.40 | 3.7 | [M-H]- | 5 | MS/MS | $C_{25}H_{50}NO_7P$ | LC-MS (-) | 0.15 | - | 36.1 | 0.097 | - | 1.6 | 13.7 | 0.29 | - | 1.4 | -16.5 | 0.36 | - | 1.2 |
| LysoPE(P-18:0/0:0) | 465.3233 | 4.68 | 2.2 | [M+H]+ | 3 | MS/MS | $C_{23}H_{48}NO_6P$ | LC-MS (+) | 0.085 | - | 33.9 | 0.037 | - | 1.5 | 13.9 | 0.30 | - | 1.1 | -15.0 | 0.21 | - | 1.1 |
| LysoPE(P-18:0/0:0) | 465.3247 | 4.85 | 3.9 | [M-H]- | 6 | MS/MS | $C_{23}H_{48}NO_6P$ | LC-MS (-) | 0.10 | - | 39.6 | 0.053 | - | 1.7 | 16.6 | 0.26 | - | 1.5 | -16.4 | 0.25 | - | 1.2 |
| PE(38:7) | 761.5006 | 12.78 | 3.6 | [M-H]- | 1 | DB | $C_{43}H_{72}NO_8P$ | LC-MS (-) | 0.020 | - | 16.0 | 0.10 | - | 1.1 | -7.7 | 0.27 | - | 1.1 | -20.4 | 0.013 | - | 1.3 |
| PE(22:6/22:6) | 835.5164 | 13.17 | 4.3 | [M-H]- | 2 | MS/MS | $C_{49}H_{74}NO_8P$ | LC-MS (-) | 0.074 | - | 25.1 | 0.057 | - | 1.3 | 4.2 | 0.64 | - | 0.8 | -16.7 | 0.093 | - | 1.1 |
| PE(32:1) | 689.5040 | 13.18 | 4.1 | [M+H]+ | 6 | DB | $C_{37}H_{72}NO_8P$ | LC-MS (+) | 0.066 | - | 21.0 | 0.11 | - | 1.2 | -5.1 | 0.63 | - | 0.9 | -21.6 | 0.033 | - | 1.3 |
| PE(20:4/18:2) | 763.5192 | 13.23 | 4.1 | [M+H]+ | 5 | MS/MS | $C_{43}H_{74}NO_8P$ | LC-MS (+) | 0.067 | - | 22.4 | 0.13 | - | 1.3 | -7.5 | 0.58 | - | 1.0 | -24.4 | 0.017 | - | 1.4 |
| PE(42:10) | 811.5167 | 13.33 | 4.7 | [M-H]- | 2 | DB | $C_{47}H_{74}NO_8P$ | LC-MS (-) | 0.012 | - | 22.9 | 0.012 | - | 1.3 | 3.7 | 0.60 | - | 0.8 | -15.7 | 0.023 | - | 1.2 |

| Compound | Mass | RT | CV | Adduct | Error | ID source | Formula | Analytical platform | p value ANOVA | pBH | % change | p value t test | p BH | VIP | % change | p value t test | p BH | VIP | % change | p value t test | p BH | VIP |
|---|---|---|---|---|---|---|---|---|---|---|---|---|---|---|---|---|---|---|---|---|---|---|
| | | | | | | | | | | | 2.5h *vs* 35 min | | | | 5.5h *vs* 35 min | | | | 5.5h *vs* 2.5h | | | |
| PE(39:9) | 855.5766 | 13.60 | 4.8 | [M+H]+ | 1 | DB | $C_{50}H_{82}NO_8P$ | LC-MS (+) | 0.063 | - | 21.6 | 0.052 | - | 1.2 | 2.4 | 0.79 | - | 0.8 | -15.8 | 0.060 | - | 1.1 |
| PE(16:0/22:6) | 763.5166 | 13.68 | 5.4 | [M+H]+ | 2 | MS/MS | $C_{43}H_{74}NO_8P$ | LC-MS (+) | 0.037 | - | 20.5 | 0.10 | - | 1.2 | -9.5 | 0.40 | - | 1.1 | -24.9 | 0.014 | - | 1.4 |
| PE(34:2) | 715.5163 | 13.73 | 5.3 | [M+H]+ | 1 | DB | $C_{39}H_{74}NO_8P$ | LC-MS (+) | 0.050 | - | 17.2 | 0.21 | - | 1.2 | -12.6 | 0.29 | - | 1.1 | -25.4 | 0.012 | - | 1.4 |
| PE(16:0/20:4) | 739.5168 | 13.85 | 3.4 | [M+H]+ | 2 | MS/MS | $C_{41}H_{74}NO_8P$ | LC-MS (+) | 0.053 | - | 22.4 | 0.085 | - | 1.3 | -3.5 | 0.73 | - | 0.9 | -21.2 | 0.026 | - | 1.2 |
| PE(32:0) | 691.5161 | 14.28 | 3.1 | [M+H]+ | 1 | DB | $C_{37}H_{74}NO_8P$ | LC-MS (+) | 0.016 | - | 20.5 | 0.091 | - | 1.2 | -11.3 | 0.27 | - | 1.1 | -26.4 | 0.0050 | - | 1.5 |
| PE(18:1/20:4) | 765.5322 | 14.38 | 3.9 | [M+H]+ | 2 | MS/MS | $C_{43}H_{76}NO_8P$ | LC-MS (+) | 0.048 | - | 23.5 | 0.11 | - | 1.3 | -7.1 | 0.55 | - | 0.9 | -24.8 | 0.018 | - | 1.4 |
| PE(44:10) | 839.5477 | 14.40 | 9.4 | [M+H]+ | 1 | DB | $C_{49}H_{78}NO_8P$ | LC-MS (+) | 0.034 | - | 12.6 | 0.30 | - | 0.8 | -15.0 | 0.10 | - | 1.2 | -24.6 | 0.014 | - | 1.3 |
| PE(16:0/18:1) | 717.5318 | 14.85 | 3.7 | [M+H]+ | 1 | MS/MS | $C_{39}H_{76}NO_8P$ | LC-MS (+) | 0.040 | - | 16.0 | 0.20 | - | 1.1 | -12.7 | 0.21 | - | 1.2 | -24.7 | 0.013 | - | 1.4 |
| PE(16:0/22:4) | 767.5556 | 15.12 | 4.7 | [M+H]+ | 12 | MS/MS | $C_{43}H_{78}NO_8P$ | LC-MS (+) | 0.069 | - | 17.8 | 0.17 | - | 1.2 | -7.9 | 0.45 | - | 0.9 | -21.9 | 0.022 | - | 1.3 |
| PE(38:4) | 789.5330 | 15.27 | 6.0 | [M+Na]+ | 6 | DB | $C_{43}H_{78}NO_8P$ | LC-MS (+) | 0.018 | - | 15.3 | 0.082 | - | 1.1 | -8.2 | 0.34 | - | 0.9 | -20.3 | 0.0036 | - | 1.3 |
| PE(18:0/18:1) | 745.5622 | 16.48 | 1.3 | [M+H]+ | 0 | MS/MS | $C_{41}H_{80}NO_8P$ | LC-MS (+) | 0.067 | - | 17.8 | 0.15 | - | 1.1 | -6.7 | 0.49 | - | 0.9 | -20.8 | 0.026 | - | 1.2 |
| PE(18:0/22:4) | 795.5781 | 16.70 | 1.9 | [M+H]+ | 0 | MS/MS | $C_{45}H_{82}NO_8P$ | LC-MS (+) | 0.055 | - | 23.0 | 0.13 | - | 1.3 | -9.0 | 0.47 | - | 1.0 | -26.0 | 0.020 | - | 1.4 |
| PE-NMe2(13D5/13M5) | 985.6507 | 17.37 | 4.8 | [M+Cl]- | 3 | DB | $C_{54}H_{96}NO_{10}P$ | LC-MS (-) | 0.056 | - | 16.4 | 0.14 | - | 1.2 | -7.6 | 0.49 | - | 1.1 | -20.6 | 0.0074 | - | 1.4 |
| PE(P-16:0/22:6) | 747.5254 | 14.20 | 2.4 | [M+H]+ | 7 | MS/MS | $C_{43}H_{74}NO_7P$ | LC-MS (+) | 0.045 | - | 18.4 | 0.088 | - | 1.2 | -5.2 | 0.57 | - | 0.9 | -20.0 | 0.019 | - | 1.2 |
| PE(O-16:0/20:4) | 725.5367 | 14.97 | 7.2 | [M+H]+ | 1 | MS/MS | $C_{41}H_{76}NO_7P$ | LC-MS (+) | 0.15 | - | 16.9 | 0.25 | - | 1.1 | -7.8 | 0.53 | - | 1.0 | -21.1 | 0.049 | - | 1.2 |
| PE(P-18:1/22:6) | 773.5370 | 15.10 | 1.5 | [M+H]+ | 1 | MS/MS | $C_{45}H_{76}NO_7P$ | LC-MS (+) | 0.047 | - | 20.1 | 0.10 | - | 1.2 | -6.9 | 0.51 | - | 1.0 | -22.5 | 0.020 | - | 1.3 |
| PE(P-18:1/16:0) | 701.5361 | 15.83 | 3.6 | [M+H]+ | 0 | MS/MS | $C_{39}H_{76}NO_7P$ | LC-MS (+) | 0.089 | - | 18.8 | 0.15 | - | 1.2 | -6.0 | 0.56 | - | 0.9 | -20.9 | 0.040 | - | 1.2 |
| PE(22:4/P-16:0) | 751.5518 | 16.05 | 4.6 | [M+H]+ | 0 | MS/MS | $C_{43}H_{78}NO_7P$ | LC-MS (+) | 0.070 | - | 18.2 | 0.20 | - | 1.2 | -10.6 | 0.34 | - | 1.1 | -24.4 | 0.025 | - | 1.3 |
| PE(P-18:1/22:4) | 777.5691 | 16.46 | 2.3 | [M+H]+ | 2 | MS/MS | $C_{45}H_{80}NO_7P$ | LC-MS (+) | 0.067 | - | 19.9 | 0.13 | - | 1.2 | -6.3 | 0.56 | - | 0.9 | -21.8 | 0.028 | - | 1.3 |
| PE(P-18:0/22:6) | 775.5566 | 16.85 | 3.9 | [M-H]- | 7 | MS/MS | $C_{45}H_{78}NO_7P$ | LC-MS (-) | 6.7X10$^{-4}$ | - | -10.3 | 0.082 | - | 0.2 | -21.1 | 4.8X10$^{-4}$ | - | 2.3 | -12.0 | 0.020 | - | 0.9 |
| PE(P-18:0/16:0) | 703.5524 | 16.89 | 4.4 | [M+H]+ | 1 | MS/MS | $C_{39}H_{78}NO_7P$ | LC-MS (+) | 0.044 | - | 13.4 | 0.23 | - | 1.1 | -11.8 | 0.24 | - | 1.1 | -22.3 | 0.0093 | - | 1.3 |
| PE(P-18:0/22:4) | 779.5833 | 17.55 | 4.5 | [M+H]+ | 0 | MS/MS | $C_{45}H_{82}NO_7P$ | LC-MS (+) | 0.083 | - | 20.8 | 0.15 | - | 1.2 | -7.5 | 0.54 | - | 1.0 | -23.5 | 0.032 | - | 1.3 |
| PE(P-45:1) / PE(O-45:0) | 855.7167 | 19.92 | 9.0 | [M-H]- | 10 | DB | $C_{50}H_{98}NO_7P$ | LC-MS (-) | 0.46 | - | -8.5 | 0.65 | - | 0.7 | -21.0 | 0.25 | - | 1.6 | -13.6 | 0.51 | - | 1.2 |
| PE(P-32:0) / PE(O-32:1) | 675.5205 | 15.32 | 2.2 | [M+H]+ | 0 | DB | $C_{37}H_{74}NO_7P$ | LC-MS (+) | 0.018 | - | 13.6 | 0.17 | - | 1.0 | -13.6 | 0.16 | - | 1.3 | -23.9 | 4.2X10$^{-3}$ | - | 1.4 |
| PE(P-37:4) / PE(O-37:5) | 737.5379 | 15.63 | 4.5 | [M+H]+ | 3 | DB | $C_{42}H_{76}NO_7P$ | LC-MS (+) | 0.047 | - | 15.8 | 0.16 | - | 1.1 | -9.3 | 0.37 | - | 1.0 | -21.7 | 0.010 | - | 1.3 |
| PE(P-40:5) / PE(O-40:6) | 777.5689 | 16.47 | 4.8 | [M+H]+ | 2 | DB | $C_{45}H_{80}NO_7P$ | LC-MS (+) | 2.9X10$^{-6}$ | pBH | -75.1 | 0.017 | pBH | 4.1 | -7.1 | 0.48 | - | 0.9 | 272.9 | 0.020 | pBH | 3.5 |
| PE(P-42:4) / PE(O-42:5) | 807.6153 | 18.65 | 6.3 | [M+H]+ | 1 | DB | $C_{47}H_{86}NO_7P$ | LC-MS (+) | 0.066 | - | 17.6 | 0.15 | - | 1.1 | -7.5 | 0.48 | - | 0.9 | -21.4 | 0.019 | - | 1.3 |
| PE(P-42:1) / PE(O-42:2) | 813.6633 | 21.15 | 1.4 | [M+H]+ | 3 | DB | $C_{47}H_{92}NO_7P$ | LC-MS (+) | 0.0035 | - | 2.9 | 0.84 | - | 0.9 | -36.3 | 0.018 | - | 2.4 | -38.1 | 0.018 | pBH | 1.9 |
| *Glycerophosphoglycerols* | | | | | | | | | | | | | | | | | | | | | | |
| LysoPG(20:1) | 538.3373 | 2.93 | 4.0 | [M-H]- | 19 | DB | $C_{26}H_{51}O_9P$ | LC-MS (-) | 0.14 | - | 39.2 | 0.095 | - | 1.7 | 10.5 | 0.48 | - | 1.4 | -20.6 | 0.20 | - | 1.3 |
| PG(38:0) | 842.5753 | 13.35 | 4.8 | [M+Cl]- | 6 | DB | $C_{44}H_{87}O_{10}P$ | LC-MS (-) | 0.013 | - | 39.6 | 0.039 | - | 1.8 | -7.5 | 0.56 | - | 1.0 | -33.7 | 0.0051 | - | 1.9 |
| PGP(44:11) | 994.4947 | 13.76 | 5.4 | [M+FA-H]- | 3 | DB | $C_{50}H_{78}O_{13}P_2$ | LC-MS (-) | 0.018 | - | 22.3 | 0.0086 | - | 1.2 | 10.4 | 0.12 | - | 1.1 | -9.7 | 0.14 | - | 0.9 |
| PG(P-16:0/13:0) | 700.4394 | 15.60 | 28.7 | [M+Cl]- | 8 | MS/MS | $C_{35}H_{69}O_9P$ | LC-MS (-) | 0.010 | - | -33.7 | 0.010 | - | 2.1 | -6.1 | 0.61 | - | 1.0 | 41.6 | 0.0064 | - | 1.8 |
| PG(P-42:1) / PG(O-42:2) | 844.6405 | 16.40 | 4.1 | [M-H]- | 18 | DB | $C_{48}H_{93}O_9P$ | LC-MS (-) | 0.12 | - | 21.2 | 0.13 | - | 1.2 | -1.5 | 0.87 | - | 0.8 | -18.7 | 0.091 | - | 1.2 |
| *Glycerophosphoinositols* | | | | | | | | | | | | | | | | | | | | | | |
| LysoPI(0:0/20:4) | 620.2986 | 2.13 | 3.4 | [M-H]- | 4 | DB | $C_{29}H_{49}O_{12}P$ | LC-MS (-) | 0.060 | - | 28.8 | 0.051 | - | 1.5 | 2.3 | 0.85 | - | 1.0 | -20.6 | 0.057 | - | 1.3 |
| LysoPI(20:4/0:0) | 620.2974 | 2.29 | 3.5 | [M-H]- | 2 | DB | $C_{29}H_{49}O_{12}P$ | LC-MS (-) | 0.16 | - | 31.3 | 0.15 | - | 1.6 | -2.2 | 0.89 | - | 1.2 | -25.5 | 0.11 | - | 1.5 |
| LysoPI(16:0) | 572.2972 | 2.55 | 4.8 | [M-H]- | 2 | DB | $C_{25}H_{49}O_{12}P$ | LC-MS (-) | 0.028 | - | 49.9 | 0.016 | - | 1.8 | 24.1 | 0.095 | - | 1.9 | -17.3 | 0.19 | - | 1.2 |
| LysoPI(20:5) | 686.2630 | 2.75 | 5.1 | [M-H+HCOONa]- | 8 | DB | $C_{29}H_{47}O_{12}P$ | LC-MS (-) | 0.20 | - | 20.9 | 0.094 | - | 1.3 | 6.0 | 0.59 | - | 1.0 | -12.3 | 0.24 | - | 1.1 |
| LysoPI(18:1) | 598.3113 | 2.87 | 8.9 | [M-H]- | 1 | DB | $C_{27}H_{51}O_{12}P$ | LC-MS (-) | 0.026 | - | 36.1 | 0.012 | - | 1.7 | 8.1 | 0.52 | - | 1.2 | -20.6 | 0.043 | - | 1.4 |
| LysoPI(18:0) | 600.3286 | 3.70 | 4.1 | [M-H]- | 2 | MS/MS | $C_{27}H_{53}O_{12}P$ | LC-MS (-) | 0.031 | - | 51.2 | 0.016 | - | 1.8 | 21.1 | 0.17 | - | 1.6 | -19.9 | 0.15 | - | 1.3 |
| LysoPI(14:0) | 580.2363 | 4.03 | 5.5 | [M-H]- | 9 | DB | $C_{23}H_{45}O_{12}P$ | LC-MS (-) | 0.070 | - | 31.6 | 0.035 | - | 1.5 | 8.7 | 0.46 | - | 1.1 | -17.4 | 0.13 | - | 1.3 |
| PI(40:10) | 902.4900 | 11.46 | 3.7 | [M+H]+ | 5 | DB | $C_{49}H_{75}O_{13}P$ | LC-MS (+) | 0.0090 | - | 28.9 | 0.0049 | - | 1.4 | 6.8 | 0.43 | - | 0.8 | -17.1 | 0.033 | - | 1.1 |
| PI(38:5) | 884.5416 | 12.00 | 1.3 | [M+H]+ | 0 | DB | $C_{47}H_{81}O_{13}P$ | LC-MS (+) | 0.036 | - | 30.7 | 0.016 | - | 1.5 | 10.4 | 0.37 | - | 1.1 | -15.6 | 0.090 | - | 1.1 |
| PI(33:2) | 866.5098 | 12.30 | 6.7 | [M+FA-H]- | 7 | DB | $C_{42}H_{77}O_{13}P$ | LC-MS (-) | 0.11 | - | 36.9 | 0.056 | - | 1.6 | 12.4 | 0.39 | - | 1.3 | -17.9 | 0.19 | - | 1.3 |
| PI(16:0/14:0) | 782.4871 | 12.75 | 4.4 | [M-H]- | 9 | MS/MS | $C_{39}H_{75}O_{13}P$ | LC-MS (-) | 0.075 | - | 22.8 | 0.074 | - | 1.2 | 1.7 | 0.85 | - | 0.8 | -17.2 | 0.077 | - | 1.1 |
| PI(40:8) | 906.5258 | 13.05 | 3.8 | [M-H]- | 0 | DB | $C_{49}H_{79}O_{13}P$ | LC-MS (-) | 0.078 | - | 17.6 | 0.20 | - | 1.3 | -9.7 | 0.40 | - | 1.2 | -23.3 | 0.023 | - | 1.5 |
| PI(20:2/20:4) | 910.5572 | 13.06 | 4.0 | [M+H]+ | 0 | MS/MS | $C_{49}H_{83}O_{13}P$ | LC-MS (+) | 0.029 | - | 39.2 | 0.023 | - | 1.6 | 12.8 | 0.28 | - | 1.2 | -19.0 | 0.13 | - | 1.1 |
| PI(18:0/20:4) | 886.5589 | 13.18 | 3.0 | [M+H]+ | 2 | MS/MS | $C_{47}H_{83}O_{13}P$ | LC-MS (+) | 0.057 | - | 25.0 | 0.036 | - | 1.3 | 3.0 | 0.79 | - | 0.9 | -17.6 | 0.052 | - | 1.2 |
| PI(20:4/16:0) | 858.5282 | 13.75 | 4.2 | [M-H]- | 3 | MS/MS | $C_{45}H_{79}O_{13}P$ | LC-MS (-) | 0.017 | - | 38.9 | 0.0095 | - | 1.6 | 16.2 | 0.13 | - | 1.4 | -16.3 | 0.11 | - | 1.2 |
| PI(38:7) | 926.5134 | 13.75 | 4.9 | [M+FA-H]- | 2 | DB | $C_{47}H_{77}O_{13}P$ | LC-MS (-) | 0.051 | - | 25.3 | 0.027 | - | 1.3 | 7.6 | 0.41 | - | 0.9 | -14.2 | 0.11 | - | 1.1 |
| PI(18:2/20:4) | 899.5629 | 15.48 | 3.3 | [M+NH4]+ | 12 | MS/MS | $C_{47}H_{79}O_{13}P$ | LC-MS (+) | 0.061 | - | -12.3 | 0.18 | - | 1.0 | 8.2 | 0.37 | - | 1.0 | 23.4 | 0.015 | - | 1.2 |
| PI(40:6) | 910.5542 | 15.50 | 6.0 | [M-H]- | 3 | DB | $C_{49}H_{83}O_{13}P$ | LC-MS (-) | 0.047 | - | 33.7 | 0.042 | - | 1.5 | 10.4 | 0.26 | - | 1.0 | -17.4 | 0.11 | - | 1.1 |
| PI(20:4/18:0) | 886.5613 | 15.64 | 4.1 | [M-H]- | 5 | MS/MS | $C_{47}H_{83}O_{13}P$ | LC-MS (-) | 0.037 | - | 21.9 | 0.026 | - | 1.3 | 4.6 | 0.54 | - | 0.7 | -14.2 | 0.066 | - | 1.1 |
| PI(O-42:0) | 1004.6791 | 4.02 | 4.0 | [M-H+HCOONa]- | 12 | DB | $C_{51}H_{101}O_{12}P$ | LC-MS (-) | 0.079 | - | 70.7 | 0.040 | - | 2.1 | 25.8 | 0.24 | - | 1.9 | -26.3 | 0.44 | - | 1.6 |
| *Glycerophosphoserines* | | | | | | | | | | | | | | | | | | | | | | |
| LysoPS(19:0) | 539.3242 | 2.16 | 4.3 | [M-H]- | 4 | DB | $C_{25}H_{50}NO_9P$ | LC-MS (-) | 0.11 | - | 33.8 | 0.089 | - | 1.5 | 6.6 | 0.61 | - | 1.1 | -20.4 | 0.29 | - | 1.3 |
| LysoPS(22:6) | 569.2778 | 2.20 | 4.4 | [M-H]- | 4 | DB | $C_{28}H_{44}NO_9P$ | LC-MS (-) | 0.069 | - | 24.1 | 0.10 | - | 1.4 | -4.8 | 0.68 | - | 1.1 | -23.3 | 0.038 | - | 1.5 |
| LysoPS(18:1) | 523.2924 | 2.76 | 6.3 | [M-H]- | 3 | DB | $C_{24}H_{46}NO_9P$ | LC-MS (-) | 0.17 | - | 23.3 | 0.11 | - | 1.2 | 6.0 | 0.54 | - | 1.0 | -14.0 | 0.21 | - | 1.0 |
| LysoPS(18:2) | 521.2756 | 2.78 | 3.6 | [M-H]- | 1 | DB | $C_{24}H_{44}NO_9P$ | LC-MS (-) | 0.062 | - | 29.7 | 0.034 | - | 1.5 | 8.2 | 0.46 | - | 1.1 | -16.6 | 0.12 | - | 1.2 |
| LysoPS(0:0/18:0) | 525.3086 | 3.53 | 4.1 | [M-H]- | 4 | MS/MS | $C_{24}H_{48}NO_9P$ | LC-MS (-) | 0.074 | - | 53.1 | 0.034 | - | 1.9 | 32.2 | 0.090 | - | 2.1 | -13.6 | 0.40 | - | 1.2 |

| Compound | Mass | RT | CV | Adduct | Error | ID source | Formula | Analytical platform | p value ANOVA | pBH | % change | p value t test | p BH | VIP | % change | p value t test | p BH | VIP | % change | p value t test | p BH | VIP |
|---|---|---|---|---|---|---|---|---|---|---|---|---|---|---|---|---|---|---|---|---|---|---|
| | | | | | | | | | | | 2.5h *vs* 35 min | | | | 5.5h *vs* 35 min | | | | 5.5h *vs* 2.5h | | | |
| LysoPS(18:0/0:0) | 525.3078 | 2.98 | 2.6 | [M+H]+ | 2 | MS/MS | $C_{24}H_{48}NO_9P$ | LC-MS (+) | 0.044 | - | 43.4 | 0.019 | - | 1.7 | 22.9 | 0.13 | - | 1.5 | -14.3 | 0.25 | - | 1.1 |
| LysoPS(18:0/0:0) | 525.3090 | 3.86 | 4.0 | [M-H]- | 5 | MS/MS | $C_{24}H_{48}NO_9P$ | LC-MS (-) | 0.028 | - | 54.4 | 0.015 | - | 2.0 | 27.8 | 0.086 | - | 2.0 | -17.2 | 0.21 | - | 1.3 |
| LysoPS(21:0/0:0) | 567.3563 | 3.10 | 3.6 | [M-H]- | 5 | DB | $C_{27}H_{54}NO_9P$ | LC-MS (-) | 0.12 | - | 37.8 | 0.071 | - | 1.7 | 10.5 | 0.47 | - | 1.3 | -19.8 | 0.19 | - | 1.3 |
| LysoPS(20:2) | 531.2962 | 3.73 | 3.4 | [M-H-H2O]- | 1 | DB | $C_{26}H_{48}NO_9P$ | LC-MS (-) | 0.079 | - | 33.9 | 0.041 | - | 1.6 | 12.2 | 0.32 | - | 1.3 | -16.1 | 0.18 | - | 1.2 |
| LysoPS(20:3) | 547.2913 | 3.87 | 7.9 | [M-H]- | 1 | DB | $C_{26}H_{46}NO_9P$ | LC-MS (-) | 0.016 | - | 51.1 | 0.0054 | - | 1.8 | 30.0 | 0.048 | - | 2.0 | -14.0 | 0.26 | - | 1.2 |
| PS(32:2) | 917.6327 | 9.78 | 5.5 | [M+FA-H]- | 3 | DB | $C_{48}H_{90}NO_{10}P$ | LC-MS (-) | 0.053 | - | 23.2 | 0.084 | - | 1.4 | -3.6 | 0.72 | - | 1.0 | -21.7 | 0.033 | - | 1.4 |
| PS(16:0/22:6) | 807.5058 | 11.50 | 2.4 | [M+H]+ | 1 | MS/MS | $C_{44}H_{74}NO_{10}P$ | LC-MS (+) | 0.10 | - | 20.2 | 0.044 | - | 1.2 | 10.2 | 0.25 | - | 1.0 | -8.3 | 0.29 | - | 0.8 |
| PS(22:4/22:6) | 883.5367 | 12.21 | 3.9 | [M+H]+ | 0 | MS/MS | $C_{50}H_{78}NO_{10}P$ | LC-MS (+) | 0.025 | - | 12.1 | 0.28 | - | 1.0 | -15.0 | 0.11 | - | 1.3 | -24.2 | 0.0059 | - | 1.4 |
| PS(22:6/17:1) | 821.5327 | 12.38 | 4.2 | [M-H]- | 12 | MS/MS | $C_{45}H_{76}NO_{10}P$ | LC-MS (-) | 0.050 | - | 17.7 | 0.11 | - | 1.2 | -6.1 | 0.46 | - | 0.9 | -20.3 | 0.031 | - | 1.3 |
| PS(39:4) | 825.5529 | 12.87 | 4.0 | [M-H]- | 1 | DB | $C_{45}H_{80}NO_{10}P$ | LC-MS (-) | 0.026 | - | 17.2 | 0.085 | - | 1.2 | -6.4 | 0.56 | - | 1.1 | -20.1 | 0.010 | - | 1.4 |
| PS(42:10) | 855.5056 | 13.00 | 4.2 | [M-H]- | 1 | DB | $C_{48}H_{74}NO_{10}P$ | LC-MS (-) | 0.024 | - | 20.0 | 0.052 | - | 1.3 | -4.5 | 0.63 | - | 0.9 | -20.5 | 0.013 | - | 1.4 |
| PS(18:1/18:1) | 787.5375 | 13.18 | 2.1 | [M+H]+ | 1 | MS/MS | $C_{42}H_{78}NO_{10}P$ | LC-MS (+) | 0.084 | - | 16.9 | 0.15 | - | 1.1 | -6.9 | 0.52 | - | 1.0 | -20.4 | 0.031 | - | 1.3 |
| PS(18:0/22:6) | 835.5373 | 13.22 | 0.8 | [M+H]+ | 1 | MS/MS | $C_{46}H_{78}NO_{10}P$ | LC-MS (+) | 0.048 | - | 18.0 | 0.080 | - | 1.2 | -6.0 | 0.56 | - | 1.0 | -20.3 | 0.019 | - | 1.3 |
| PS(18:0/20:4) | 811.5389 | 13.38 | 4.4 | [M+H]+ | 3 | MS/MS | $C_{44}H_{78}NO_{10}P$ | LC-MS (+) | 0.046 | - | 25.2 | 0.087 | - | 1.3 | -4.2 | 0.70 | - | 0.9 | -23.5 | 0.022 | - | 1.3 |
| PS(35:0) | 823.5498 | 13.65 | 5.4 | [M+FA-H]- | 10 | DB | $C_{41}H_{80}NO_{10}P$ | LC-MS (-) | 0.069 | - | 21.4 | 0.082 | - | 1.2 | -2.2 | 0.80 | - | 0.8 | -19.4 | 0.040 | - | 1.3 |
| PS(38:2) | 837.5430 | 13.66 | 7.2 | [M+Na]+ | 8 | DB | $C_{44}H_{82}NO_{10}P$ | LC-MS (+) | 0.024 | - | 20.6 | 0.058 | - | 1.2 | -7.1 | 0.48 | - | 1.0 | -22.9 | 0.011 | - | 1.3 |
| PS(18:1/18:0) | 789.5463 | 14.31 | 2.4 | [M+H]+ | 7 | MS/MS | $C_{42}H_{80}NO_{10}P$ | LC-MS (+) | 0.071 | - | 19.7 | 0.11 | - | 1.2 | -4.6 | 0.64 | - | 0.9 | -20.3 | 0.035 | - | 1.2 |
| PS(40:4) | 839.5663 | 14.57 | 3.9 | [M+H]+ | 2 | MS/MS | $C_{46}H_{82}NO_{10}P$ | LC-MS (+) | 0.047 | - | 17.4 | 0.13 | - | 1.1 | -8.6 | 0.41 | - | 1.0 | -22.2 | 0.014 | - | 1.3 |
| PS(20:4/22:6) | 891.4777 | 15.70 | 4.3 | [M+Cl]- | 5 | MS/MS | $C_{48}H_{74}NO_{10}P$ | LC-MS (-) | 0.036 | - | 20.4 | 0.039 | - | 1.4 | -5.1 | 0.73 | - | 0.8 | -21.2 | 0.0061 | - | 1.4 |
| PS(38:1) | 817.5830 | 15.72 | 3.2 | [M+H]+ | 0 | DB | $C_{44}H_{84}NO_{10}P$ | LC-MS (+) | 0.069 | - | 21.7 | 0.11 | - | 1.2 | -3.4 | 0.72 | - | 0.8 | -20.6 | 0.041 | - | 1.2 |
| PS(41:5) | 887.5344 | 16.35 | 4.7 | [M+Cl]- | 12 | DB | $C_{47}H_{82}NO_{10}P$ | LC-MS (-) | 0.0024 | - | 24.3 | 0.0047 | - | 1.2 | -0.1 | 0.99 | - | 0.5 | -19.6 | 0.0011 | - | 1.4 |
| PS(40:1) | 845.6159 | 17.36 | 3.9 | [M+H]+ | 1 | DB | $C_{46}H_{88}NO_{10}P$ | LC-MS (+) | 0.091 | - | 22.0 | 0.082 | - | 1.2 | 1.7 | 0.86 | - | 0.8 | -16.7 | 0.065 | - | 1.1 |
| PS(46:5) | 957.6247 | 17.37 | 4.7 | [M+Cl]- | 2 | DB | $C_{52}H_{92}NO_{10}P$ | LC-MS (-) | 0.044 | - | 15.6 | 0.13 | - | 1.2 | -8.3 | 0.42 | - | 1.1 | -20.6 | 0.0075 | - | 1.4 |
| PS(46:4) | 969.6533 | 17.84 | 3.7 | [M+FA-H]- | 15 | DB | $C_{52}H_{94}NO_{10}P$ | LC-MS (-) | 0.017 | - | 20.2 | 0.0089 | - | 1.3 | 5.4 | 0.44 | - | 1.0 | -12.4 | 0.042 | - | 1.0 |
| *Glycerophosphoglycerophosphoglycerols* | | | | | | | | | | | | | | | | | | | | | | |
| CL(88:5) | 1539.1222 | 13.70 | 5.6 | [M+H]+ | 6 | DB | $C_{87}H_{160}O_{17}P_2$ | LC-MS (+) | 0.050 | - | 27.0 | 0.11 | - | 1.5 | -15.2 | 0.36 | - | 1.3 | -33.2 | 0.014 | - | 1.7 |
| CL(78:4) | 1563.1234 | 13.73 | 5.3 | [M+Na]+ | 8 | DB | $C_{87}H_{162}O_{17}P_2$ | LC-MS (+) | 0.035 | - | 25.1 | 0.12 | - | 1.4 | -12.2 | 0.37 | - | 1.2 | -29.8 | 0.011 | - | 1.6 |
| CL(79:2) | 1581.1527 | 14.05 | 1.1 | [M+Na]+ | 3 | DB | $C_{88}H_{168}O_{17}P_2$ | LC-MS (+) | 0.079 | - | 22.1 | 0.064 | - | 1.3 | 1.5 | 0.88 | - | 0.8 | -16.9 | 0.060 | - | 1.1 |
| CL(75:2) | 1520.1313 | 14.15 | 3.6 | [M+NH4]+ | 6 | DB | $C_{84}H_{160}O_{17}P_2$ | LC-MS (+) | 0.048 | - | 17.0 | 0.10 | - | 1.2 | -6.6 | 0.52 | - | 0.9 | -20.2 | 0.010 | - | 1.3 |
| CL(76:2) | 1517.1354 | 14.20 | 3.8 | [M+H]+ | 4 | DB | $C_{85}H_{162}O_{17}P_2$ | LC-MS (+) | 0.045 | - | 15.4 | 0.17 | - | 1.1 | -9.9 | 0.32 | - | 1.1 | -21.9 | 0.011 | - | 1.3 |
| CL(78:4) | 1541.1391 | 14.20 | 1.5 | [M+H]+ | 7 | DB | $C_{87}H_{162}O_{17}P_2$ | LC-MS (+) | 0.034 | - | 23.7 | 0.12 | - | 1.3 | -11.4 | 0.37 | - | 1.2 | -28.4 | 0.011 | - | 1.5 |
| CL(76:2) | 1555.0955 | 14.35 | 3.5 | [M+K]+ | 7 | DB | $C_{85}H_{162}O_{17}P_2$ | LC-MS (+) | 0.045 | - | 25.2 | 0.062 | - | 1.3 | -4.5 | 0.68 | - | 0.9 | -23.8 | 0.023 | - | 1.3 |
| CL(76:1) | 1519.1602 | 14.75 | 3.1 | [M+H]+ | 10 | DB | $C_{85}H_{164}O_{17}P_2$ | LC-MS (+) | 0.054 | - | 22.2 | 0.11 | - | 1.3 | -7.7 | 0.52 | - | 1.0 | -24.5 | 0.022 | - | 1.4 |
| CL(75:2) | 1541.0840 | 15.35 | 3.4 | [M+K]+ | 10 | DB | $C_{82}H_{160}O_{17}P_2$ | LC-MS (+) | 0.038 | - | 22.0 | 0.087 | - | 1.3 | -7.8 | 0.50 | - | 1.0 | -24.5 | 0.014 | - | 1.4 |
| CL(76:4) | 1535.0984 | 15.40 | 2.3 | [M+Na]+ | 13 | DB | $C_{85}H_{158}O_{17}P_2$ | LC-MS (+) | 0.042 | - | 17.1 | 0.16 | - | 1.1 | -10.6 | 0.31 | - | 1.1 | -23.7 | 0.014 | - | 1.4 |
| CL(80:2) | 1595.1768 | 15.45 | 5.4 | [M+Na]+ | 2 | DB | $C_{89}H_{170}O_{17}P_2$ | LC-MS (+) | 0.042 | - | 20.0 | 0.16 | - | 1.2 | -12.5 | 0.32 | - | 1.1 | -27.1 | 0.010 | - | 1.5 |
| CL(80:8) | 1561.0964 | 15.57 | 3.9 | [M+H]+ | 1 | DB | $C_{89}H_{158}O_{17}P_2$ | LC-MS (+) | 0.030 | - | 18.1 | 0.18 | - | 1.2 | -14.7 | 0.22 | - | 1.3 | -27.8 | 0.0064 | - | 1.5 |
| CL(73:0) | 1517.0834 | 15.58 | 2.5 | [M+K]+ | 10 | DB | $C_{82}H_{160}O_{17}P_2$ | LC-MS (+) | 0.043 | - | 25.5 | 0.12 | - | 1.4 | -11.4 | 0.42 | - | 1.2 | -29.4 | 0.013 | - | 1.6 |
| CL(75:2) | 1503.1056 | 16.28 | 2.8 | [M+H]+ | 5 | DB | $C_{84}H_{160}O_{17}P_2$ | LC-MS (+) | 0.080 | - | 21.9 | 0.13 | - | 1.3 | -6.5 | 0.60 | - | 1.0 | -23.3 | 0.035 | - | 1.4 |
| CL(77:2) | 1569.1109 | 16.28 | 4.1 | [M+K]+ | 7 | DB | $C_{86}H_{164}O_{17}P_2$ | LC-MS (+) | 0.039 | - | 15.2 | 0.13 | - | 1.1 | -9.2 | 0.36 | - | 1.0 | -21.2 | 0.0074 | - | 1.3 |
| CL(75:0) | 1529.1204 | 16.46 | 2.3 | [M+Na]+ | 4 | DB | $C_{84}H_{164}O_{17}P_2$ | LC-MS (+) | 0.051 | - | 24.5 | 0.12 | - | 1.3 | -8.8 | 0.50 | - | 1.1 | -26.8 | 0.019 | - | 1.5 |
| **Hydroxy acids and derivatives** | | | | | | | | | | | | | | | | | | | | | | |
| Glyceric acid | 106.0266 | 10.61 | 8.8 | - | - | Fiehn | $C_3H_6O_4$ | GC-MS | 0.0053 | pBH | 38.6 | 0.024 | pBH | 1.5 | 43.2 | 0.0053 | pBH | 1.3 | 3.3 | 0.68 | - | 0.6 |
| Lactic acid | 90.0317 | 6.78 | 3.3 | - | - | Fiehn | $C_3H_6O_3$ | GC-MS | 0.080 | - | -4.6 | 0.28 | - | 0.9 | -33.5 | 0.025 | pBH | 1.3 | -30.3 | 0.026 | - | 1.2 |
| Malic acid | 134.0215 | 12.67 | 6.1 | - | - | Fiehn | $C_4H_6O_5$ | GC-MS | 0.035 | - | 34.9 | 0.027 | - | 1.3 | 0.8 | 0.95 | - | 0.8 | -25.3 | 0.042 | - | 1.2 |
| **Non-metal oxoanionic compounds** | | | | | | | | | | | | | | | | | | | | | | |
| Phosphoric acid | 97.9769 | 8.454 | 8.0 | - | - | Fiehn | $H_3PO_4$ | GC-MS | 0.059 | - | 28.3 | 0.026 | - | 1.2 | 8.7 | 0.38 | - | 0.7 | -15.3 | 0.15 | - | 0.9 |
| **Organic phosphoric acids and derivatives** | | | | | | | | | | | | | | | | | | | | | | |
| O-Phosphoethanolamine | 141.0191 | 16.112 | 21.5 | - | - | Fiehn | $C_2H_8NO_4P$ | GC-MS | 0.014 | - | 0.4 | 0.98 | - | 0.6 | -37.7 | 0.036 | pBH | 1.3 | -37.9 | 0.013 | - | 1.3 |
| **Organooxygen compounds** | | | | | | | | | | | | | | | | | | | | | | |
| 2,3-butanediol | 90.0681 | 6.543 | 3.3 | - | - | Fiehn | $C_4H_{10}O_2$ | GC-MS | 0.082 | - | -4.6 | 0.28 | - | 0.9 | -33.4 | 0.025 | pBH | 1.3 | -30.2 | 0.026 | - | 1.2 |
| 2,3-diphosphoglycerate | 265.9592 | 20.308 | 12.1 | - | - | Fiehn | $C_3H_8O_{10}P_2$ | GC-MS | 0.018 | - | 42.9 | 0.026 | pBH | 1.6 | 11.2 | 0.88 | - | 0.6 | -22.1 | 0.052 | - | 1.1 |
| Threonic acid | 136.0372 | 13.545 | 7.8 | - | - | Fiehn | $C_4H_8O_5$ | GC-MS | $6.5X10^{-5}$ | pBH | 58.7 | 0.0013 | pBH | 1.9 | 77.3 | $1.9X10^{-4}$ | pBH | 1.7 | 11.7 | 0.16 | - | 0.7 |
| **Sphingolipids [SP]** | | | | | | | | | | | | | | | | | | | | | | |
| *Acidic glycosphingolipids* | | | | | | | | | | | | | | | | | | | | | | |
| NeuAcα2-3Galβ-Cer(d18:1/16:0) | 1036.6696 | 2.75 | 3.8 | [M+FA-H]- | 4 | DB | $C_{51}H_{94}N_2O_{16}$ | LC-MS (-) | 0.077 | - | 83.9 | 0.041 | - | 2.3 | 26.3 | 0.28 | - | 1.9 | -31.3 | 0.40 | - | 1.7 |
| (3'-sulfo)Galbeta-Cer(d36:1) | 807.5549 | 11.81 | 6.0 | [M+H]+ | 2 | DB | $C_{42}H_{81}NO_{11}S$ | LC-MS (+) | 0.051 | - | 27.0 | 0.024 | - | 1.3 | 8.4 | 0.38 | - | 0.9 | -14.7 | 0.10 | - | 1.0 |
| Ganglioside GM1 (18:1/18:0) | 1545.8733 | 13.14 | 3.6 | [M-H]- | 2 | DB | $C_{73}H_{131}N_3O_{31}$ | LC-MS (-) | 0.059 | - | 21.0 | 0.048 | - | 1.3 | -0.4 | 0.97 | - | 0.9 | -17.7 | 0.047 | - | 1.3 |
| KDNalpha2-3Galbeta1-3GalNAcbeta1-4(KDNalpha2-3)Galbeta1-4Glcbeta-Cer(d18:1/24:0) | 1838.9937 | 13.40 | 4.2 | [M-2H]-2 | 8 | DB | $C_{86}H_{154}N_2O_{39}$ | LC-MS (-) | 0.023 | - | 24.4 | 0.022 | - | 1.3 | -0.5 | 0.96 | - | 0.7 | -20.0 | 0.025 | - | 1.4 |

| Compound | Mass | RT | CV | Adduct | Error | ID source | Formula | Analytical platform | p value ANOVA | pBH | % change | p value t test | p BH | VIP | % change | p value t test | p BH | VIP | % change | p value t test | p BH | VIP |
|---|---|---|---|---|---|---|---|---|---|---|---|---|---|---|---|---|---|---|---|---|---|---|
| | | | | | | | | | | | 2.5h *vs* 35 min | | | | 5.5h *vs* 35 min | | | | 5.5h *vs* 2.5h | | | |
| Ganglioside GM3 (d36:1) | 1180.7411 | 13.52 | 3.8 | [M-H]- | 3 | DB | $C_{59}H_{108}N_2O_{21}$ | LC-MS (-) | 0.070 | - | 14.6 | 0.23 | - | 1.1 | -9.3 | 0.37 | - | 1.0 | -20.9 | 0.013 | - | 1.3 |
| (3'-sulfo)Galbeta-Cer(d40:1) | 863.6140 | 15.20 | 4.6 | [M+H]+ | 2 | DB | $C_{46}H_{89}NO_{11}S$ | LC-MS (+) | 0.070 | - | 18.3 | 0.13 | - | 1.1 | -6.0 | 0.53 | - | 0.9 | -20.5 | 0.033 | - | 1.2 |
| (3'-sulfo)Galbeta-Cer(d18:1/20:0(2OH)) | 851.5771 | 15.63 | 5.0 | [M-H]- | 2 | DB | $C_{44}H_{85}NO_{12}S$ | LC-MS (-) | 0.081 | - | 21.1 | 0.41 | - | 1.1 | -3.9 | 0.54 | - | 0.8 | -20.6 | 0.10 | - | 1.2 |
| (3'-sulfo)Galbeta-Cer(d42:1) | 891.6430 | 16.68 | 4.0 | [M+H]+ | 5 | DB | $C_{48}H_{93}NO_{11}S$ | LC-MS (+) | 0.085 | - | 20.4 | 0.12 | - | 1.2 | -3.8 | 0.80 | - | 0.8 | -20.1 | 0.035 | - | 1.2 |
| *Ceramides* | | | | | | | | | | | | | | | | | | | | | | |
| Cer(d18:2/18:0) | 563.5312 | 14.45 | 4.8 | [M-H]- | 6 | DB | $C_{36}H_{69}NO_3$ | LC-MS (-) | 0.11 | - | 23.7 | 0.11 | - | 1.3 | -2.9 | 0.81 | - | 1.0 | -21.5 | 0.075 | - | 1.4 |
| Cer(d38:2) | 637.5650 | 16.22 | 4.0 | [M+FA-H]- | 1 | DB | $C_{38}H_{73}NO_3$ | LC-MS (-) | 0.15 | - | 22.8 | 0.11 | - | 1.1 | -0.1 | 1.00 | - | 0.9 | -18.6 | 0.12 | - | 1.2 |
| Cer(d18:1/22:1) | 619.5820 | 17.28 | 2.5 | [M+H]+ | 14 | MS/MS | $C_{40}H_{77}NO_3$ | LC-MS (+) | 0.022 | - | 21.2 | 0.017 | - | 1.1 | 27.4 | 0.011 | - | 1.6 | 5.1 | 0.57 | - | 0.8 |
| Cer(d34:0(2OH)) | 588.5495 | 17.38 | 3.1 | [M+NH4]+ | 9 | DB | $C_{34}H_{69}NO_5$ | LC-MS (+) | 0.041 | - | 25.9 | 0.043 | - | 1.3 | 0.3 | 0.98 | - | 0.8 | -20.3 | 0.038 | - | 1.2 |
| Cer(d18:1/24:1) | 647.6223 | 17.57 | 3.5 | [M+H]+ | 1 | DB | $C_{42}H_{81}NO_3$ | LC-MS (+) | 0.092 | - | 21.6 | 0.13 | - | 1.2 | -4.1 | 0.70 | - | 0.9 | -21.2 | 0.048 | - | 1.2 |
| Cer(d40:0) | 691.6124 | 18.01 | 4.3 | [M-H+HCOONa]- | 5 | DB | $C_{40}H_{81}NO_3$ | LC-MS (-) | 0.18 | - | 22.2 | 0.056 | - | 1.1 | 10.0 | 0.37 | - | 1.0 | -10.0 | 0.35 | - | 1.0 |
| *Neutral glycosphingolipids* | | | | | | | | | | | | | | | | | | | | | | |
| GlcCer(d38:1) / GalCer(d38:1) | 755.6265 | 13.55 | 4.1 | [M+H]+ | 1 | DB | $C_{44}H_{85}NO_8$ | LC-MS (+) | 0.068 | - | 19.3 | 0.13 | - | 1.1 | -5.2 | 0.57 | - | 0.8 | -20.5 | 0.037 | - | 1.2 |
| GlcCer(d18:1/24:0) / GalCer(d18:1/24:0) | 811.6862 | 16.68 | 3.6 | [M+H]+ | 5 | MS/MS | $C_{48}H_{93}NO_8$ | LC-MS (+) | 0.073 | - | 21.5 | 0.13 | - | 1.2 | -5.7 | 0.58 | - | 0.9 | -22.4 | 0.039 | - | 1.3 |
| GlcCer(41:2) / GalCer(41:2) | 817.6411 | 17.21 | 4.8 | [M+Na]+ | 0 | DB | $C_{47}H_{89}NO_8$ | LC-MS (+) | 0.0050 | - | 23.2 | 0.025 | - | 1.2 | -9.1 | 0.35 | - | 1.0 | -26.3 | 0.0026 | - | 1.4 |
| GlcCer(d18:1/24:1) / GalCer(d18:1/24:1) | 855.6811 | 18.22 | 4.8 | [M+FA-H]- | 2 | DB | $C_{48}H_{91}NO_8$ | LC-MS (-) | 0.044 | - | 20.4 | 0.066 | - | 1.2 | 0.1 | 0.99 | - | 0.6 | -16.9 | 0.049 | - | 1.1 |
| GlcCer(d38:2) | 805.5978 | 18.40 | 5.9 | [M+Cl]- | 19 | DB | $C_{44}H_{83}NO_9$ | LC-MS (-) | 0.013 | - | 10.5 | 0.17 | - | 0.9 | -12.7 | 0.060 | - | 1.5 | -21.0 | 0.0073 | - | 1.5 |
| *Phosphonosphingolipids* | | | | | | | | | | | | | | | | | | | | | | |
| N-palmitoyl-D-sphingosyl-1-(2-aminoethyl)phosphonate | 666.5088 | 18.99 | 6.7 | [M+Na]+ | 2 | DB | $C_{36}H_{73}N_2O_5P$ | LC-MS (+) | 0.022 | pBH | 19.2 | 0.0019 | - | 1.2 | 28.5 | 0.0035 | - | 1.9 | 7.8 | 0.089 | - | 0.6 |
| *Phosphosphingolipids* | | | | | | | | | | | | | | | | | | | | | | |
| SM(d34:1) | 702.5687 | 12.32 | 2.7 | [M+H]+ | 1 | MS/MS | $C_{39}H_{79}N_2O_6P$ | LC-MS (+) | 0.017 | - | 28.4 | 0.067 | - | 1.5 | -9.2 | 0.48 | - | 1.0 | -29.3 | 0.0038 | - | 1.5 |
| SM(d16:1/18:0) | 748.5745 | 12.67 | 3.7 | [M+FA-H]- | 2 | MS/MS | $C_{39}H_{79}N_2O_6P$ | LC-MS (-) | 0.015 | - | 24.8 | 0.060 | - | 1.4 | -7.5 | 0.48 | - | 1.1 | -25.8 | 0.0042 | - | 1.6 |
| SM(d36:2) | 728.5848 | 13.01 | 3.3 | [M+H]+ | 2 | DB | $C_{41}H_{81}N_2O_6P$ | LC-MS (+) | 0.018 | - | 38.3 | 0.065 | - | 1.5 | -16.3 | 0.33 | - | 1.2 | -39.5 | 0.0048 | - | 1.8 |
| SM(34:0) | 704.5819 | 13.03 | 8.3 | [M+H]+ | 2 | DB | $C_{39}H_{81}N_2O_6P$ | LC-MS (+) | 0.010 | - | 28.3 | 0.048 | - | 1.4 | -10.6 | 0.40 | - | 1.0 | -30.4 | 0.0033 | - | 1.6 |
| SM(d38:0) | 650.5293 | 13.21 | 6.0 | [M+H]+ | 11 | DB | $C_{35}H_{75}N_2O_6P$ | LC-MS (+) | 0.035 | - | 21.5 | 0.092 | - | 1.3 | -9.3 | 0.44 | - | 1.1 | -25.3 | 0.011 | - | 1.4 |
| SM(18:1/18:1) | 774.5900 | 13.35 | 4.2 | [M+FA-H]- | 2 | MS/MS | $C_{41}H_{81}N_2O_6P$ | LC-MS (-) | 0.020 | - | 41.2 | 0.046 | - | 1.8 | -10.6 | 0.34 | - | 1.2 | -36.7 | 0.023 | - | 1.9 |
| SM(d38:2) | 756.6095 | 14.82 | 4.6 | [M+H]+ | 7 | DB | $C_{43}H_{85}N_2O_6P$ | LC-MS (+) | 0.034 | - | 42.7 | 0.062 | - | 1.6 | -8.5 | 0.94 | - | 0.9 | -35.9 | 0.016 | - | 1.6 |
| SM(d30:0) | 650.5289 | 15.40 | 6.8 | [M+H]+ | 11 | DB | $C_{35}H_{75}N_2O_6P$ | LC-MS (+) | 0.032 | - | 15.9 | 0.14 | - | 1.1 | -11.0 | 0.29 | - | 1.1 | -23.2 | 0.0078 | - | 1.4 |
| SM(d38:1) | 758.6294 | 15.78 | 1.6 | [M+H]+ | 1 | MS/MS | $C_{43}H_{87}N_2O_6P$ | LC-MS (+) | 0.034 | - | 22.3 | 0.083 | - | 1.3 | -6.4 | 0.55 | - | 0.9 | -23.4 | 0.013 | - | 1.3 |
| SM(40:1) | 786.6632 | 17.52 | 3.8 | [M+H]+ | 2 | MS/MS | $C_{45}H_{91}N_2O_6P$ | LC-MS (+) | 0.055 | - | 25.9 | 0.095 | - | 1.3 | -5.1 | 0.69 | - | 0.9 | -24.6 | 0.023 | - | 1.3 |
| *Sphingoid bases* | | | | | | | | | | | | | | | | | | | | | | |
| Sphingosine | 299.2833 | 1.86 | 0.9 | [M+H]+ | 3 | MS/MS | $C_{18}H_{37}NO_2$ | LC-MS (+) | 0.021 | pBH | 0.0 | 1.00 | - | 0.7 | -26.3 | 0.025 | pBH | 2.2 | -26.3 | $6.3X10^{-4}$ | - | 1.51 |
| **Sterol Lipids [ST]** | | | | | | | | | | | | | | | | | | | | | | |
| *Steroid conjugates* | | | | | | | | | | | | | | | | | | | | | | |
| (25R)-3alpha,7alpha-dihydroxy-5beta-cholestan-27-oyl taurine | 609.3279 | 2.75 | 3.7 | [M-H+HCOONa]- | 6 | DB | $C_{29}H_{51}NO_6S$ | LC-MS (-) | 0.11 | - | 30.8 | 0.060 | - | 1.5 | 10.5 | 0.39 | - | 1.2 | -15.5 | 0.21 | - | 1.2 |
| Taurocholic acid | 515.2856 | 9.78 | 7.6 | [M-H]- | 12 | DB | $C_{26}H_{45}NO_7S$ | LC-MS (-) | 0.0084 | - | 20.7 | 0.018 | - | 1.3 | -2.5 | 0.72 | - | 0.7 | -19.2 | 0.017 | - | 1.3 |
| *Sterols* | | | | | | | | | | | | | | | | | | | | | | |
| CE(20:4) | 694.5677 | 24.08 | 8.9 | [M+Na]+ | 2 | DB | $C_{47}H_{76}O_2$ | LC-MS (+) | $2.4X10^{-4}$ | pBH | -14.1 | 0.15 | - | 1.0 | -50.0 | 0.0043 | pBH | 3.5 | -41.9 | 0.0067 | pBH | 2.1 |
| **Prenol Lipids** | | | | | | | | | | | | | | | | | | | | | | |
| *Isoprenoids* | | | | | | | | | | | | | | | | | | | | | | |
| Retinol | 286.2309 | 5.48 | 1.0 | [M+H]+ | 4 | DB | $C_{20}H_{30}O$ | LC-MS (+) | 0.0057 | - | 15.2 | 0.21 | - | 1.1 | -21.8 | 0.058 | - | 1.6 | -32.1 | $8.9X10^{-4}$ | - | 1.7 |
| **Pyridinecarboxylic acids and derivatives** | | | | | | | | | | | | | | | | | | | | | | |
| Nicotinamide | 122.0480 | 12.628 | 5.3 | - | - | Fiehn | $C_6H_6N_2O$ | GC-MS | 0.019 | - | 11.2 | 0.21 | - | 0.8 | -12.6 | 0.13 | - | 0.8 | -21.3 | 0.040 | pBH | 1.1 |
| Ubiquinol-10 | 886.6841 | 22.93 | 5.4 | [M+Na]+ | 3 | DB | $C_{59}H_{92}O_4$ | LC-MS (+) | 0.045 | pBH | 2.2 | 0.86 | - | 0.7 | -35.8 | 0.010 | - | 2.4 | -37.2 | 0.013 | pBH | 1.9 |
| **Pyridines and derivatives** | | | | | | | | | | | | | | | | | | | | | | |
| Niacinamide | 122.0483 | 0.78 | 3.5 | [M+H]+ | 2 | MS/MS | $C_6H_6N_2O$ | LC-MS (+) | 0.025 | - | 6.6 | 0.46 | - | 0.8 | -15.7 | 0.074 | - | 1.3 | -20.9 | 0.0054 | - | 1.2 |
| **Tricarboxylic acids and derivatives** | | | | | | | | | | | | | | | | | | | | | | |
| Citric acid | 192.0270 | 16.484 | 4.2 | - | - | Fiehn | $C_6H_8O_7$ | GC-MS | 0.0053 | pBH | 0.1 | 0.99 | - | 0.7 | -40.1 | 0.036 | pBH | 1.5 | -40.2 | 0.0019 | pBH | 1.5 |
| **Multiple candidates** | | | | | | | | | | | | | | | | | | | | | | |
| 2-oxo-5-amino-pentanoic acid / Trans-3-hydroxy-L-proline | 131.0589 | 0.75 | 5.1 | [M-H]- | 6 | DB | $C_5H_9NO_3$ | LC-MS (-) | 0.016 | - | 22.5 | 0.092 | - | 1.3 | -15.7 | 0.15 | - | 1.7 | -31.2 | 0.0042 | - | 1.8 |
| D-Ribose 1-diphosphate / 2-Deoxy-D-ribose 1,5-bisphosphate | 329.9643 | 1.08 | 4.1 | [M+Cl]- | 10 | MS/MS | $C_5H_{12}O_{10}P_2$ | LC-MS (-) | 0.42 | - | -27.0 | 0.34 | - | 0.8 | -29.7 | 0.27 | - | 1.6 | -3.7 | 0.89 | - | 1.2 |
| Cholesterol sulfate | 466.3134 | 6.16 | 4.3 | [M-H]- | 4 | DB | $C_{27}H_{46}O_4S$ | LC-MS (-) | 0.19 | - | 38.1 | 0.25 | - | 1.1 | 2.6 | 0.69 | - | 0.7 | -25.7 | 0.21 | - | 1.0 |
| LysoPA(20:0) | 466.3134 | 6.16 | 4.3 | [M-H]- | 16 | DB | $C_{23}H_{47}O_7P$ | LC-MS (-) | 0.19 | - | 38.1 | 0.25 | - | 1.1 | 2.6 | 0.69 | - | 0.7 | -25.7 | 0.21 | - | 1.0 |
| PI-Cer(d34:1) | 779.5232 | 12.19 | 3.9 | [M-H]- | 10 | DB | $C_{40}H_{78}NO_{11}P$ | LC-MS (-) | 0.056 | - | 21.6 | 0.075 | - | 1.2 | -1.4 | 0.87 | - | 0.8 | -18.9 | 0.045 | - | 1.2 |
| (3'-sulfo)Galbeta-Cer(d34:1) | 779.5232 | 12.19 | 3.9 | [M-H]- | 2 | DB | $C_{40}H_{77}NO_{11}S$ | LC-MS (-) | 0.056 | - | 21.6 | 0.075 | - | 1.2 | -1.4 | 0.87 | - | 0.8 | -18.9 | 0.045 | - | 1.2 |
| PE(38:6) | 779.5232 | 12.19 | 3.9 | [M-H]- | 17 | DB | $C_{43}H_{74}NO_9P$ | LC-MS (-) | 0.056 | - | 21.6 | 0.075 | - | 1.2 | -1.4 | 0.87 | - | 0.8 | -18.9 | 0.045 | - | 1.2 |
| GlcCer(d34:1) | 716.5846 | 13.25 | 3.6 | [M+NH4]+ | 10 | DB | $C_{40}H_{77}NO_8$ | LC-MS (+) | 0.041 | - | 22.7 | 0.15 | - | 1.4 | -12.6 | 0.41 | - | 1.1 | -28.8 | 0.0039 | - | 1.5 |

| Compound | Mass | RT | CV | Adduct | Error | ID source | Formula | Analytical platform | p value ANOVA | pBH | % change | p value t test | p BH | VIP | % change | p value t test | p BH | VIP | % change | p value t test | p BH | VIP |
|---|---|---|---|---|---|---|---|---|---|---|---|---|---|---|---|---|---|---|---|---|---|---|
| | | | | | | | | | | | 2.5h *vs* 35 min | | | | 5.5h *vs* 35 min | | | | 5.5h *vs* 2.5h | | | |
| SM(35:1) / PE-Cer(d38:1) | 716.5846 | 13.25 | 3.6 | [M+H]+ | 2 | DB | $C_{40}H_{81}N_2O_6P$ | LC-MS (+) | 0.041 | - | 22.7 | 0.15 | - | 1.4 | -12.6 | 0.41 | - | 1.1 | -28.8 | 0.0039 | - | 1.5 |
| PG(42:6) | 882.5263 | 13.59 | 4.9 | [M+Cl]- | 10 | DB | $C_{48}H_{79}O_{10}P$ | LC-MS (-) | 0.022 | - | 49.2 | 0.021 | - | 1.7 | 15.4 | 0.23 | - | 1.6 | -22.7 | 0.059 | - | 1.3 |
| PI(O-36:3) / PI(P-36:2) | 882.5263 | 13.59 | 4.9 | [M+Cl]- | 15 | DB | $C_{45}H_{83}O_{12}P$ | LC-MS (-) | 0.022 | - | 49.2 | 0.021 | - | 1.7 | 15.4 | 0.23 | - | 1.6 | -22.7 | 0.059 | - | 1.3 |
| SM(d37:1) / PE-Cer(d40:1) | 744.6064 | 15.00 | 4.3 | [M+H]+ | 11 | DB | $C_{42}H_{85}N_2O_6P$ | LC-MS (+) | 0.062 | - | 18.7 | 0.16 | - | 1.2 | -7.9 | 0.50 | - | 0.9 | -22.4 | 0.011 | - | 1.3 |
| 3-O-Sulfogalactosylceramide (d32:1) | 891.6511 | 19.84 | 4.1 | [M-H]- | 5 | DB | $C_{48}H_{93}NO_{11}S$ | LC-MS (-) | 0.075 | - | 20.0 | 0.060 | - | 1.2 | 3.1 | 0.68 | - | 0.9 | -14.1 | 0.086 | - | 1.0 |
| LacCer(d36:0) | 891.6511 | 19.84 | 4.1 | [M-H]- | 15 | DB | $C_{48}H_{93}NO_{13}$ | LC-MS (-) | 0.075 | - | 20.0 | 0.060 | - | 1.2 | 3.1 | 0.68 | - | 0.9 | -14.1 | 0.086 | - | 1.0 |

CV, coefficient of variation of metabolites in QC samples; Mass error, expressed in PPM; ID source, identification source, DB corresponds to database source; *p* value ANOVA, global *p* value obtained by one-way ANOVA test; pBH, global *p* value obtained with Benjamini–Hochberg correction test; % change in the specific comparison; *p* value t test, obtained by t test for each comparison; pBH, *p* value obtained with Benjamini–Hochberg correction test for each comparison; *P* values marked with "†" correspond to the corrected *p* value obtained after Benjamini-Hochberg correction test; VIP, VIP values higher than 1 were considered as significant.
